# Large-Scale Assessment of Language, Speech, and Movement in Autism and ADHD with AI

**DOI:** 10.1101/2025.10.20.682864

**Authors:** Aimar Silvan, Lucas C. Parra, Alexandre R. Franco, Adriana Di Martino, Michael P. Milham, Jens Madsen

## Abstract

Attention-Deficit/Hyperactivity Disorder (ADHD) and Autism Spectrum Disorder (ASD) frequently co-occur with overlapping traits. We investigated whether automated behavioral analysis during a clinician-child interview can identify distinct, objective features of the two conditions. Analyzing audio-video recordings of 2,529 youths (ages 5–22) in a broad community sample, multivariate models revealed that language difficulties often attributed to ADHD are primarily explained by age, cognitive ability, or co-occurring ASD. Increased motor activity specifically marked hyperactive-impulsive ADHD, but not ASD or inattentive ADHD. ASD was uniquely characterized by divergent narrative production and idiosyncratic responses, alongside a distinct vocal profile of higher pitch and dysphonia, despite structurally intact language. While these digital behavioral measures correlate with most diagnostic categories and age, the joint analysis effectively separates the effects of ASD from ADHD. These findings show that scalable digital assessment from recorded clinical interviews can disentangle overlapping ASD and ADHD diagnoses into domain-specific behavioral signatures.

## 1. Introduction

Attention-Deficit/Hyperactivity Disorder (ADHD) and Autism Spectrum Disorder (ASD) are both prevalent conditions with long-term functional impacts [1–4]. ADHD involves persistent inattention and/or hyperactivity, while ASD is characterized by challenges with social interaction and restricted, repetitive behaviors and interests [5]. These heterogeneous conditions frequently co-occur [6–12], sharing genetic factors [13–16] and converging variations in large-scale brain networks [17–22].

Current diagnostic assessments for ASD and ADHD rely on structured laboratory tasks, which can lack ecological validity [23–25], or on parent and teacher reports, which can be subject to recall or rater bias [26]. Furthermore, traditional observational research typically focuses on isolated behavioral domains and is constrained by small sample sizes due to the high cost and time required for manual behavioral coding. This has traditionally limited research to smaller samples that do not capture the vast heterogeneity inherent in ADHD and ASD in more natural settings. Meanwhile, routine clinical visits provide rich, ecologically valid opportunities to observe patients’ spontaneous behaviors in a naturalistic social context. Here, we test whether large-scale, automated analysis of naturalistic clinical interactions can disentangle overlapping ADHD and ASD diagnoses into separable behavioral dimensions.

Multiple studies have looked at behavioral presentations of ASD and ADHD. However, studying these conditions in isolation may obscure their true clinical presentation [27]. Disentangling overlapping ADHD and ASD traits into separable behavioral dimensions is conceptually and clinically critical to better understand genetic heterogeneity [15], improve precise disorder characterization [28], and ensure timely diagnosis [29].

We leveraged the Healthy Brain Network (HBN) cohort [30], a large, cross-sectional community sample of neurotypical and neurodivergent youth. The data included a video-recorded, semi-structured interview where participants were asked about an emotive animated film they just watched. This easily scalable interview was designed to elicit language, emotional expression, and social-cognitive interpretations.

While core deficits in attention, sensory processing, and executive functioning are central to these conditions, evaluating them typically requires highly structured, constrained cognitive tasks [31]. In contrast, during naturalistic social interactions, the most prominent and clinically observable manifestations of neurodivergence are how a child speaks, sounds, and moves. We therefore prioritized the automated assessment of three directly observable domains: motor activity, language use, and speech prosody. State-of-the-art AI enables the objective, simultaneous extraction of these metrics from audio-video recordings. In the following, we review relevant literature in the context of ADHD and ASD.

Effective communication depends on both structural language (vocabulary, syntax) and pragmatic language skills (social application, topic coherence) [32]. Social communication deficits are a core diagnostic feature of ASD but not ADHD [5]. However, studies using informant-based assessments suggest a significant degree of overlap, with both groups demonstrating difficulties in both pragmatic and structural language [32–34].

Beyond fundamental language skills, perspective-taking is also essential for discourse and narrative comprehension and construction [35]. This is often described as Theory of Mind (ToM), i.e., the ability to infer the mental and emotional states of others. Difficulties in perspective-taking have long been associated with ASD [36]. Studies using automated language analysis have identified repetitive speech, atypical social narratives in ASD [35,37], and narrative disorganization in ADHD [38]. However, these methods typically evaluate fewer than 50 participants [34] and analyze conditions separately, limiting generalizability and obscuring shared versus distinct traits. We address these gaps by jointly modeling structural, pragmatic, and higher-order semantic and narrative abilities across both diagnoses to develop a detailed model of traits in both ADHD and ASD on an unprecedented scale.

Alongside communication deficits, atypical prosody is a clinical hallmark of ASD [37]. Recent systematic reviews link ASD to higher mean pitch and greater pitch variability [39], though findings regarding loudness, speech rate, and voice quality remain inconsistent due to methodological heterogeneity and small sample sizes [39, 40]. Furthermore, separate reports on ADHD suggest that it also presents with distinct prosodic features [41, 42]. What is currently unclear is whether this presentation is specific to ADHD, or if they result from the co-occurring ASD, or whether they interact to exacerbate the effects. Automated vocal analysis shows diagnostic promise [43–45], but it has not yet been used to determine which vocal signatures are due to the co-occurrence, or which can be uniquely attributed to either group.

Motor differences are central to both ADHD and ASD, yet their behavioral signatures are rarely quantified under comparable conditions. While ASD has been associated with diverse motor patterns [46, 47], it remains unclear which of these reflect distinct motor phenotypes or are shared manifestations of ADHD.

Further complicating matters, language, speech, and motor coordination develop rapidly with age, and symptom presentations can shift [48]. For instance, hyperactivity in ADHD often decreases while inattention persists into adolescence [49]. Without rigorously modeling key covariates like age, sex [50, 51], and cognitive ability (IQ) [52, 53], observed markers may simply reflect typical developmental or demographic variance rather than true psychopathology.

This first analysis of the HBN semi-structured interview presents an opportunity to simultaneously observe these diverse behavioral domains at an unprecedented scale. Using clinical diagnoses, we apply a multivariate approach to identify joint drivers of behavioral variance across this cross-sectional developmental cohort. Deviating from prior work that treats co-occurring ADHD and ASD as a distinct categorical group [54–56], we model ADHD Inattention, ADHD Hyperactivity, and ASD as additive dimensions consistent with the spirit of the NIMH Research Domain Criteria [57, 58]. Treating co-occurring ASD+ADHD as a separate diagnostic category is important in clinical management of this subgroup. However, a categorical analysis artificially fragments variance and reduces statistical power. In contrast, the additive dimensional analysis has higher statistical power and can naturally account for covariates such as age and IQ. But most importantly, it isolates the unique variance attributable to each condition. By mapping digital phenotypes to these joint predictors rather than forcing participants into mutually exclusive categories, our approach isolates condition-specific behavioral signatures and helps reconcile prior inconsistencies in the literature.

## 2. Results

### 2.1. A semi-structured interview of a community sample for automatic behavioral analysis

To measure naturalistic behaviors, 2,529 participants (aged 5-22) of the HBN cohort were video-recorded during a semi-structured interview after watching “The Present”, a brief animated film in which a boy’s interaction with a three-legged puppy culminates in the revelation that both share a missing leg. During the interview, participants were asked identical questions one at a time (Table S1) probing narrative recall, factual memory, emotional interpretation, and perspective-taking. We used state-of-the-art computational tools to derive objective, quantitative metrics from the video-recorded interviews, covering expressive language, vocal prosody, as well as facial and upper body movements (Fig. 1a).

**Figure 1:**
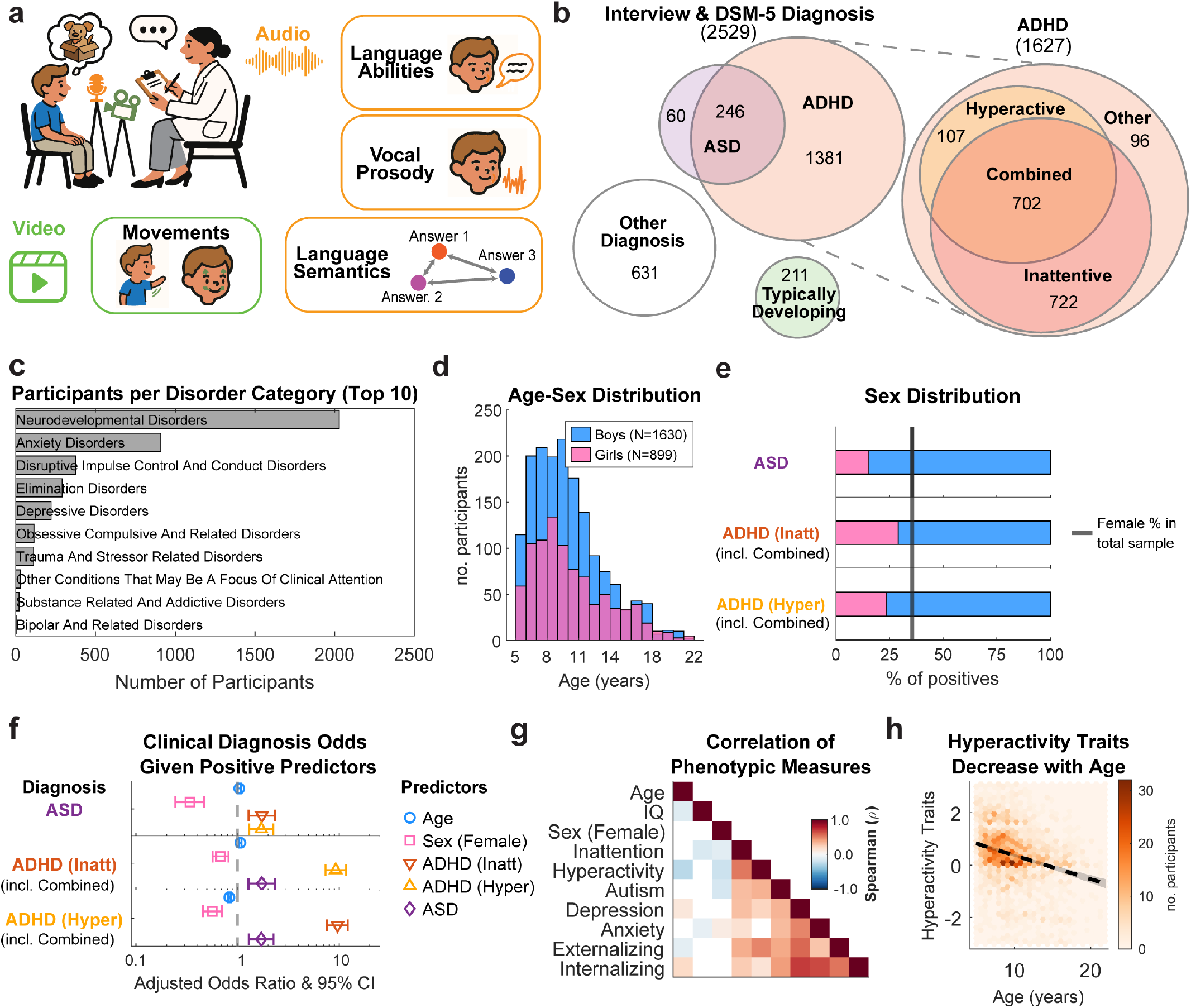
Overview of the study cohort. a) Using AI for objective behavioral characterization. Video recordings were processed to quantify upper body and facial movements, while audio was transcribed and analyzed to extract metrics of language structure, semantic content, and vocal prosody. b) Venn diagram showing the distribution and overlap of the primary diagnostic groups. c) The 12 most common clinician-assigned DSM-5 diagnostic categories in the full cohort (*N* = 2, 529), see Fig. S1 for specific co-occurring diagnoses for the ADHD and ASD children. d) Age and sex distribution of the full cohort, see Fig. S1 for age-sex distributions per diagnostic group in panel b. e) Sex distribution within the children with ASD and/or ADHD diagnostic, ADHD-Combined are here considered positive in both hyperactive and inattentive diagnostic status. The black line indicates chance occurrence based on the sex ratio in the cohort. f) Adjusted odds ratios showing the likelihood of a co-occurring positive diagnosis in individuals with positive specific predictors (or one-year change in the case of age). Females are less likely than males to receive an ADHD or ASD diagnosis (OR *<* 1). g) Spearman correlation matrix, displaying only significant associations (*p <* 0.01, Bonferroni corrected, white otherwise). The matrix illustrates that clinical instruments for specific neurodevelopmental traits, like ASSQ for autism and SWAN for ADHD, are highly correlated with other measurements like depression (MFQ), anxiety (SCARED), and internalizing/externalizing behaviors (CBCL). They also correlate with age, IQ, and sex, which complicates the attribution of observed behavior to any single instrument scale independently. h) Two-dimensional histogram showing the negative correlation between age and hyperactivity traits (SWAN scale) (*ρ*(2095) = *−*0.27, *p <* 0.001). Full names for clinical instruments (SWAN, ASSQ, MFQ, SCARED, CBCL) are provided in the ‘Clinical Instruments’ section in the Methods. Detailed model statistics for panel F are available in Supplementary Section S1.1.

The cohort was diagnostically rich and clinically complex (Fig. 1b, c & Table 1). The sample exhibited an overall male-to-female ratio of approximately 2:1 (Fig. 1d), with lower than chance prevalence in females (15.4% of the ASD group and 29-23.6% of the ADHD group; Fig. 1e). Consistent with the clinical literature [12], a clinician-confirmed diagnostic status of either ADHD-Inattentive or ADHD-Hyperactive significantly increased the odds of co-occurring ASD (Adjusted Odds Ratios (OR) = 1.734 and 1.716, respectively, Fig. 1f). Clinical overlap extended to instrument ratings, which were highly correlated and strongly influenced by age and sex (Fig. 1g). For instance, parent-rated hyperactivity traits decreased significantly with age (*ρ*(2095) = *−*0.27, *p <* 0.001, Bonferroni corrected, Fig. 1h), consistent with typical developmental trajectories [49]. The same was also reflected in the effect of age on the prevalence of ADHD-Hyperactive status (Fig. 1f, OR = 0.831). To summarize Fig. 1f, ASD, Hyper-active, Inattentive, Age and Sex are all associated with one another.

**Table 1:**
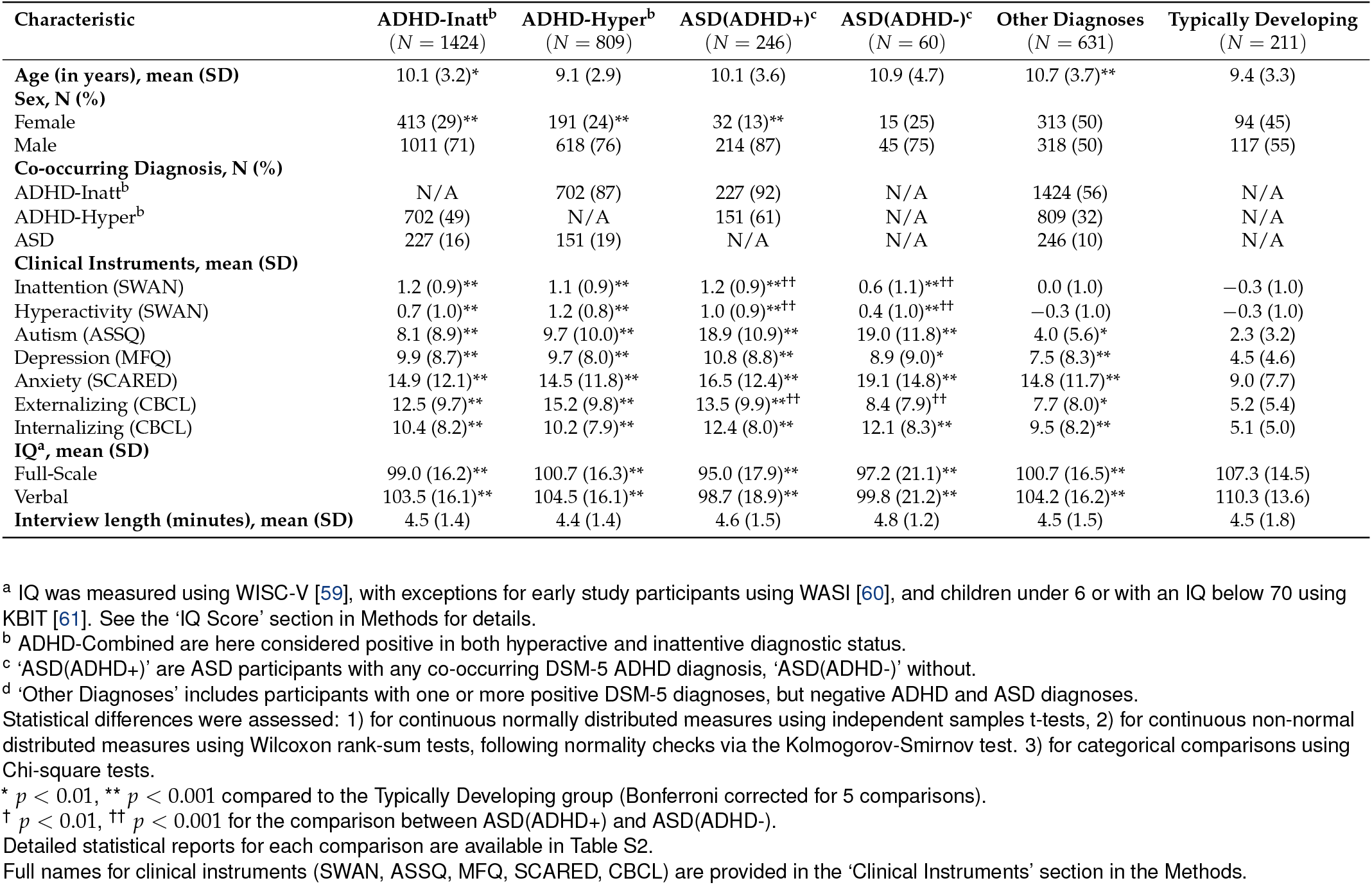
Demographic, clinical, and interview characteristics.

Note that in Fig. 1e-f and in all our analyses, ADHD-Combined presentation is coded as both Inattentive and Hyperactive. To distinguish this coding with two binary variables (Inattentive, Hyperactive) from the three DSM-5 ADHD presentations (Inattentive, Hyperactive, Combined), we will refer to these as diagnostic status and presentation, respectively.

### 2.2. Developmental factors dominate over diagnostic effects in structural language

To characterize naturalistic communication at scale, we processed the audio recordings of the interview using a state-of-the-art AI pipeline that integrates automatic transcription with Large Language Model (LLM)-based diarization. From these transcripts, we extracted objective metrics of structural and pragmatic language abilities (Fig. 2a).

**Figure 2:**
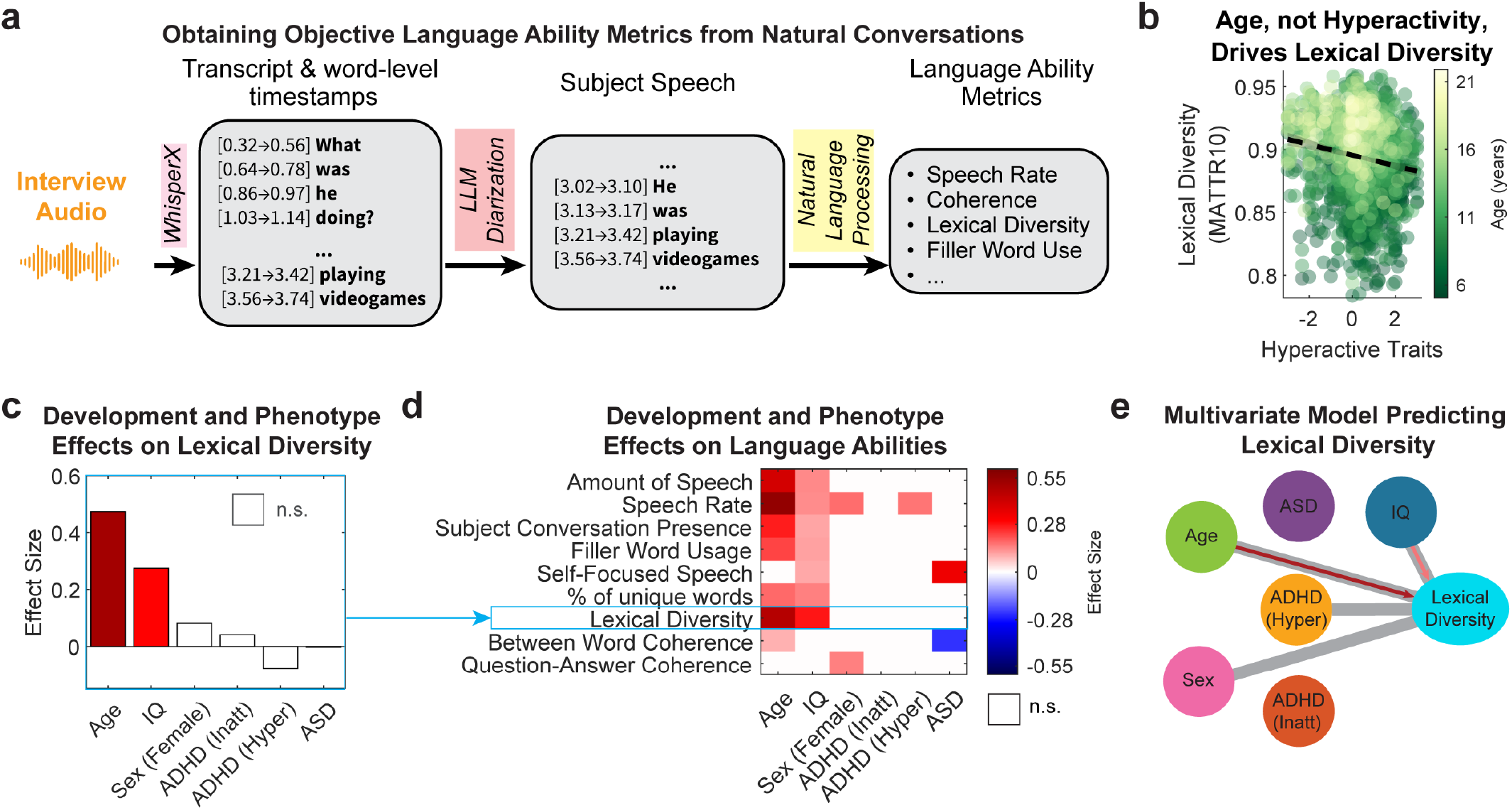
Language abilities are primarily driven by development, not by ADHD or ASD. a) The computational pipeline used to extract objective structural and pragmatic language metrics from transcribed audio. b) Hyperactive traits (SWAN) and lexical diversity (Moving-Average Type-Token-Ratio with a window of 10) are correlated (*ρ*(2049) = *−*0.16, *p <* 0.001). However, the color gradient, representing age, reveals that this association is confounded by development. c) Effect sizes from the multivariate model predicting lexical diversity. Effect sizes represent the unique contribution of each factor (Sex coded as Female = 1, Male = 0). For binary variables, the reported effects are Cohen’s *d*, while for numerical variables, we report Cohen’s *f* (see Methods). d) A summary matrix of effect sizes from the multivariate model applied to a range of language metrics. The results show the consistent and strong effects of age and IQ, and the general lack of diagnostic effects across most measured aspects of language abilities. Significance cut-off at *p <* 0.01 (white otherwise; Bonferroni-corrected across the 9 models tested). Detailed multivariate model reports are available in Supplementary Section S1.2. e) Result of the dimensional multivariate regression analysis for lexical diversity specifically. Gray links are significant associations in a pairwise correlation, but only red arrows were statistically significant after including covariates in the multivariate model. Missing links indicate no significant association. Associations shown in Fig. 1f are omitted here for simplicity.

To isolate the unique behavioral contributions of each diagnosis while accounting for high co-occurrence and developmental maturation, we employed a multivariate regression model. This approach coded diagnostic status as independent binary variables (ASD, ADHD-Inattentive, ADHD-Hyperactive) while simultaneously controlling for Age, Sex, and IQ (Fig. 2c).

This multivariate approach demonstrated that structural language abilities are overwhelmingly driven by development, not by ADHD or ASD. For example, a simple univariate analysis suggested a significant negative association between lexical diversity and hyperactivity (*ρ*(2049) = *−*0.16, *p <* 0.001, Bonferroni Corrected, Fig. 2b). However, the multivariate model revealed this variation was actually driven by the effect of Age (Effect Size=0.474; effect sizes (ES) reported in this study represent Cohen’s *d* for binary and Cohen’s *f* for continuous variables; see Methods) and IQ (ES=0.275) (Fig. 2c-d). This pattern held across almost all structural language metrics (Fig. 2e). Older participants spoke more (ES=0.404) and faster (ES=0.553), while females spoke faster (ES=0.171) and answered more coherently (ES=0.149) than males. The only significant diagnostic markers identified in basic language use were an increase in self-referential speech (first-person pronouns) (ES=0.337), and reduced between-word coherence (ES=-0.242) specific to participants with ASD, as well as faster speech rate in ADHD-Hyperactivity (ES=0.164).

Our planned analysis was to model ADHD and ASD as independent additive effects. We also explored interaction terms to determine whether the co-occurrence is stronger/weaker than the sum of their parts, but found no significant interactions. Additionally, we explored the alternative approach of modeling the co-occurrence as a separate diagnostic category (ADHD+ASD). These exploratory analyses are treated in the Methods under “Alternative analysis approaches” along with Age interactions and other post hoc analyses.

### 2.3. ASD is uniquely associated with atypical narrative and idiosyncratic responses

While structural language remained largely intact across diagnoses, impairments in social communication may often stem from higher-order difficulties in narrative construction, thematic interpretation, and social-emotional expression. To objectively quantify these functions, we computationally compared the semantic content of each participant’s interview responses to those of their age-matched, Typically Developing peers, generating an objective measure of “answer typicality” (Fig. 3a-c).

**Figure 3:**
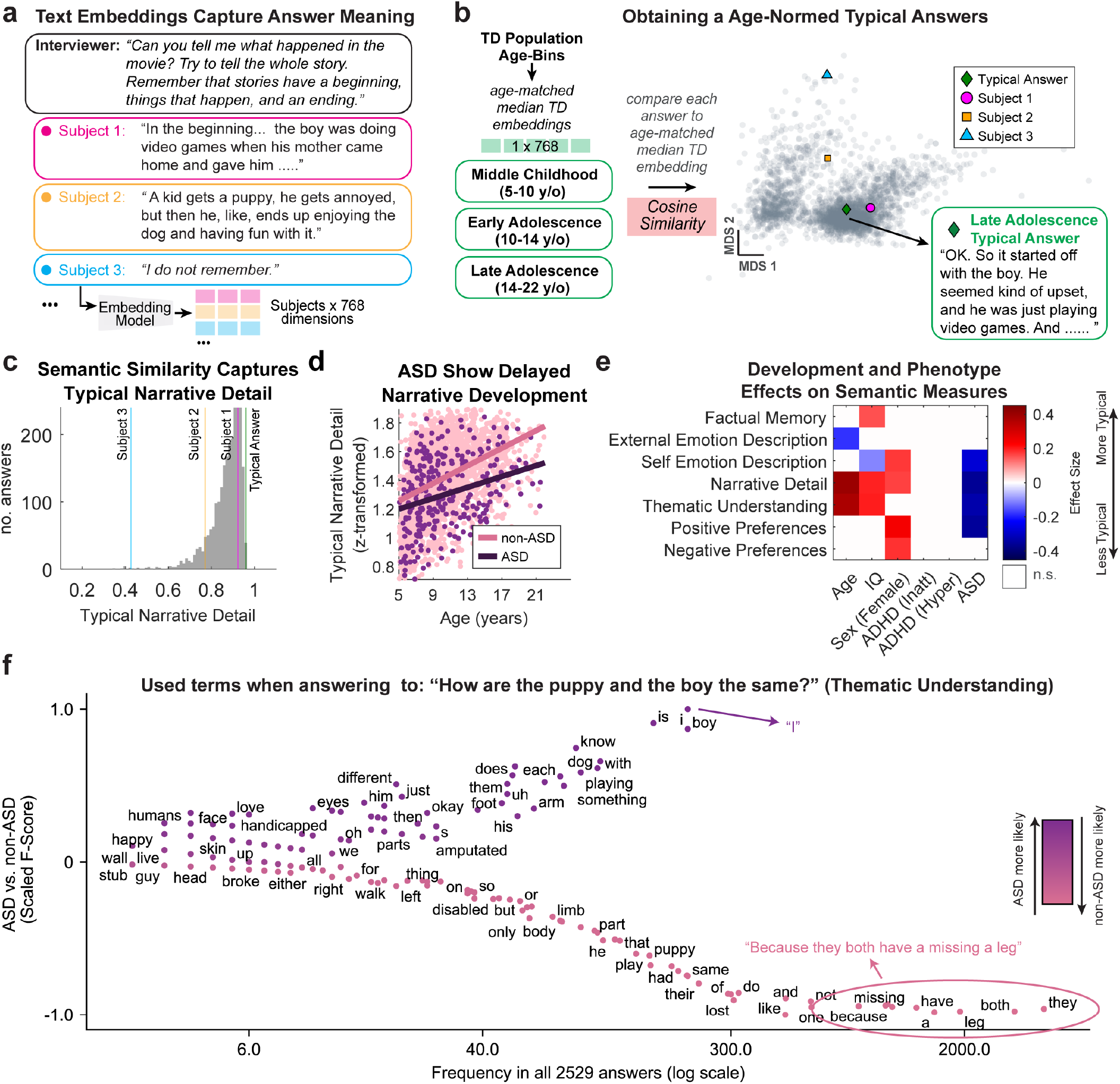
Semantic analysis reveals ASD-specific effects on narrative and semantic response patterns. a) The computational pipeline for semantic analysis. Participants’ answers to 23 predefined interview questions were parsed and converted into numerical semantic representations (embeddings). b) Multidimensional Scaling (MDS) representation of the answer embeddings, where similar embeddings are represented closer together. A “typical answer” is obtained from the semantic (cosine) similarity to the median embedding of the corresponding age-matched TD group answers. c) Semantic similarity to the age-appropriate prototype is used as a measure of ‘typicality’. For the narrative recall question in panel A, higher similarity reflects a more detailed and developmentally expected recount of the narrative relative to age-matched peers. Subject 1 provides a detailed answer (high similarity), whereas Subject 3 does not recall the story (low similarity). d) Joint distribution illustrating the positive association between age and the production of more typical narratives, but the delayed development in the ASD group compared to the non-ASD group. e) Matrix of effect sizes from the multivariate model, assessing the impact of diagnosis and demographic factors on various semantic domains probed by the interview questions (e.g., factual memory, emotional description, and thematic understanding). Only effect sizes with *p <* 0.01, after Bonferroni correction, are shown in color. See Table S1 for a full list of questions and the typical answers for each age bin, and Supplementary Section S1.3 for detailed multivariate model reports. f) Scatterplot of terms distinguishing ASD (top, purple) from non-ASD (bottom, pink) participants in responses to the question related to thematic understanding, using scattertext [62] (See Methods section ‘Visualization of Distinctive Terms’ for details, and Fig. S2 for visualization of other questions).

After controlling for Age, our models revealed a significant, divergent semantic response profile specific to ASD (Fig. 3e). An ASD diagnosis was uniquely associated with producing less typical narratives (ES=-0.358, Fig. 3d), showing different thematic understanding – such as failing to convey that the boy and the puppy were similar because they were both missing a leg (Fig. 3f) – (ES=-0.330), providing idiosyncratic descriptions of their own emotions (ES=-0.277), and expressing atypical positive preferences (ES=-0.0352). In contrast, neither the ADHD-Inattentive nor ADHD-Hyperactive statuses were significantly associated with any of these semantic and narrative domains.

Despite measuring similarity of responses relative to age-matched controls, age remained a powerful predictor across some semantic language use measures (Fig. 3e). This pattern is consistent with increasing convergence towards age-normative responses, rather than a drift in the semantics of the responses. Indeed, the exemplary ‘typical answers’ for each age bin were remarkably similar (Table S1), suggesting no age-related difficulties in understanding or answering the questions within the Typically Developing population. Consistent with social-cognitive maturation [48], older participants provided more consistent, detailed narratives (ES=0.402) and recalled the movie’s main message more typically (ES=0.373). Similarly, higher IQ was associated with more typical narrative abilities (ES=0.0194), thematic understanding (ES=0.198), and factual memory (ES=0.154). We also observed a secondary finding related to sex, with females tending to produce more typical narratives (ES=0.169), and positive and negative preferences (ES=0.242 & 0.181) compared to male participants (Fig. 3e).

### 2.4. Autism Spectrum Disorder has a distinct vocal profile

The acoustic patterns of pitch, loudness, and rhythm are critical for conveying the intent of an utterance. To this end, we employed an automated pipeline to extract a comprehensive set of acoustic features from the segmented speech of each child, including measures of pitch, loudness, and voice quality (Fig. 4a).

**Figure 4:**
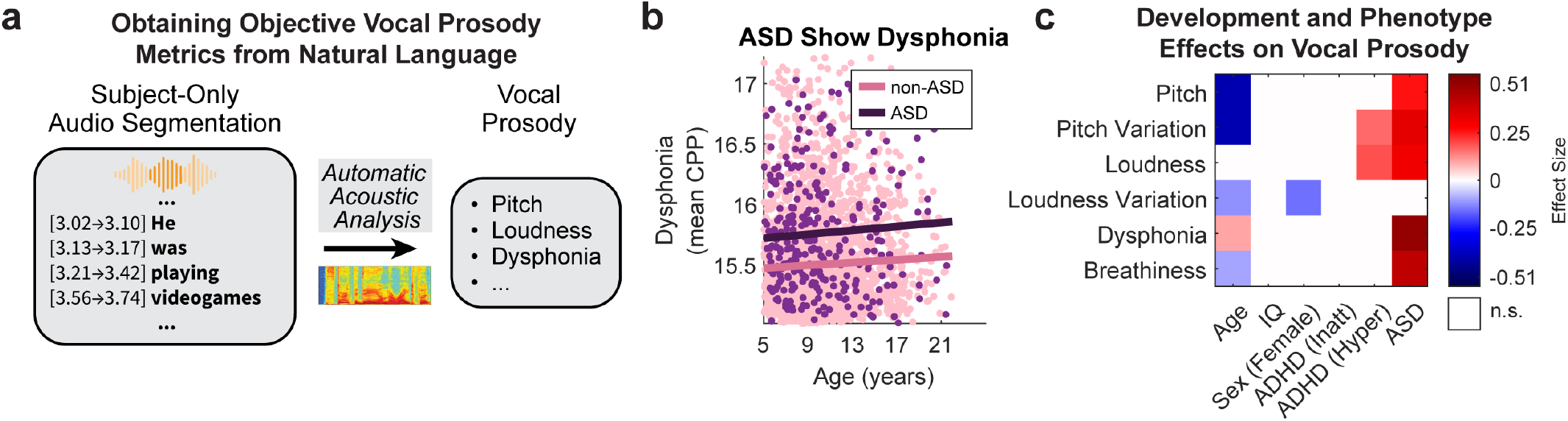
Automatic vocal prosody metrics reveal an ASD-specific acoustic profile. a) Automatic extraction of vocal prosody features involved transcribing and diarizing speech, thus isolating the children’s speech from that of the adult interviewer, and extracting audio-based features using automatic acoustic analysis [63]. b) ASD participants showed increased dysphonia compared to non-ASD participants, which may be related to an “unusual” voice in ASD. c) Effect Size matrix illustrating the effects of development and ASD on vocal prosody. Effect Sizes in this matrix are estimated again using a multivariate regression model, with *p <* 0.01, after Bonferroni correction. See Supplementary Section S1.4 for detailed multivariate model reports.

Our automated acoustic analysis revealed a distinct vocal signature specific to ASD. Controlling for covariates, ASD was significantly associated with increased mean pitch (ES=0.262), greater pitch variability (ES=0.162), and higher vocal loudness (ES=0.307) (Fig. 4c). Furthermore, participants with ASD exhibited higher dysphonia (ES=0.509, Fig. 4b) and increased breathiness (ES=0.416). ADHD-Hyperactivity was only associated with pitch variability (ES=0.162) and loudness (ES=0.191). Age again proved to be a dominant predictor, with older participants naturally exhibiting reduced pitch (ES=-0.398) and pitch variability (ES=-0.399).

### 2.5. Increased motor activity is uniquely associated with ADHD-Hyperactive diagnostic status

Motor behaviors are relevant diagnostic features of ASD and ADHD, with hyperactivity being a key diagnostic criterion in ADHD, and stereotypical and repetitive motor behaviors being among the defining criteria for ASD. Yet motor behaviors are rarely quantified objectively in naturalistic settings. Here, we applied computer vision models to extract objective frame-to-frame displacement metrics for the upper body, head, eyes, and face (Fig. 5a). Consistent with typical maturation, age was a strong negative predictor of movement categories (ES ranging from -0.419 to -0.27).

**Figure 5:**
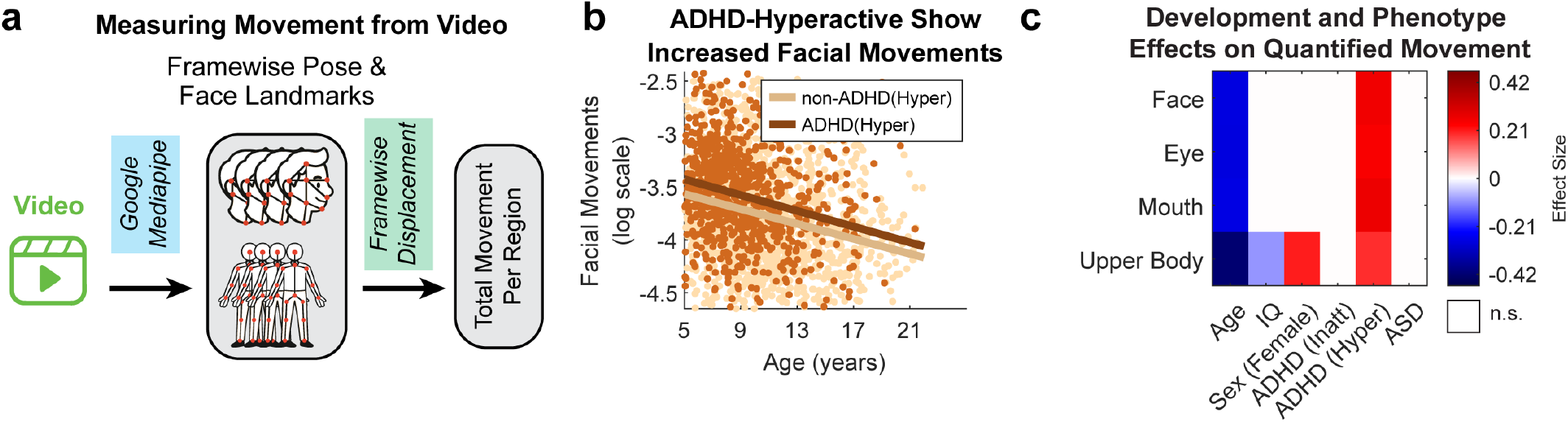
Associations of demographics and phenotype with upper body and face movements. a) We use Google’s Mediapipe computer vision models to obtain framewise locations for anatomical and facial landmarks. We then calculate the average framewise displacement for the different regions of interest. B) ADHD-Hyperactive youth show increased facial movements across the whole developmental spectrum, compared to non-ADHD Hyperactive. C) Effect Size matrix illustrating the effects of development and an ADHD-Hyperactive status on all measures of face and upper body movements. Effect Sizes in this matrix are estimated using multivariate regression, with *p <* 0.01, after Bonferroni correction. See Supplementary Section S1.5 for detailed multivariate model reports.

Once developmental decreases in movement were accounted for, we identified a highly specific motor signature: ADHD-Hyperactive status was robustly associated with increased movement across all measured upper body and facial regions (ES=0.194-0.266) (Fig. 5b-c). Conversely, neither ASD nor ADHD-Inattentive status were linked to significant changes in motor activity, confirming that objective video-derived kinematics isolate the hyperactive-impulsive domain of ADHD without generalizing to the inattentive presentation or ASD.

## 3. Discussion

This study applied automated digital phenotyping to a large community sample to isolate behavioral dimensions of ASD and ADHD, demonstrating that while narrative divergence and atypical speech uniquely characterize ASD, increased motor activity specifically marks ADHD-Hyperactivity. Crucially, differences in the structural language and narrative metrics examined here, which are often attributed to ADHD, are better explained by age, IQ, and co-occurring ASD.

Previous standardized assessments suggest ADHD involves structural and pragmatic language impairments [64, 65], with community prevalence estimated at 40-45% [66, 67]. However, standardized tests impose heavy demands on executive functions like sustained attention and working memory. Consistent with other naturalistic studies [68] our conversational analysis shows that once age, IQ, and ASD are controlled, structural, conversational language in ADHD, as captured by our automated metrics, appears intact, except for faster speech rate. However, this does not rule out potential ADHD-related difficulties in other linguistic domains not captured here, such as real-time reciprocity or complex discourse management [69].

Such results extend to higher-order semantic analysis. ADHD was not independently associated with neither narrative nor answer atypicality, indicating that previously reported differences may reflect developmental factors or diagnostic overlap rather than ADHD-specific effects [38, 70]. Apparent communication difficulties in ADHD may instead likely reflect executive dysfunction, such as poor inhibition, impacting speech organization, rather than fundamental deficits in linguistic comprehension and higher-order narrative interpretation or construction [71, 72]. Distinct semantic and narrative divergence thus remains a critical differentiator indicating co-occurring ASD in youths with ADHD [73].

Conversely, our ASD cohort exhibited mostly typical structural language trajectories, supporting classic conceptualizations of the disorder as a core social communication deficit [5]. The primary structural deviation was increased self-referential speech via first-person pronouns, reflecting an asynchrony between linguistic and social development [74], and reduced between-word coherence. Most ASD communication differences emerged strictly in higher-order domains: participants produced less typical narratives, missed overarching thematic ‘gists’ [75], and provided idiosyncratic emotion descriptions [36].

It is critical to carefully define what these semantic similarity measures do and do not capture, as the Theory of Mind (ToM) construct has faced significant recent scrutiny for lacking specificity, predictive validity, and reproducibility, with many ToM deficits actually reflecting hidden language barriers or differing cognitive styles rather than an inability to mentalize [76]. Our semantic metric captures statistical divergence from the neurotypical consensus in how a story is interpreted and recounted; it does not directly measure an underlying psychological inability to understand others. However, because this semantic divergence in ASD persists even after controlling for clinically measured language abilities (CELF-Screener [77]; see Fig. S3), it cannot be dismissed as a simple vocabulary deficit. Instead, this naturalistic validation aligns with computational analyses suggesting that autistic individuals employ distinct, idiosyncratic narrative styles and words (Fig. 3d, Fig. S2) to describe social and emotive contexts [78, 79], reflecting neurodivergent thematic prioritization rather than a fundamental language deficit. Alternatively, they may reflect difficulties in narrative construction or social understanding, memory, or attentional factors [80].

Using automated acoustic analysis, we objectively quantified the long-observed atypical prosody of ASD [40]. Our analysis confirmed higher pitch and pitch variability [39], while clarifying past inconsistencies by identifying increased dysphonia and breathiness as distinct, ADHD-independent ASD markers, validating clinical descriptions of “hoarse” voices [81]. Except for pitch variability and loudness, these effects were absent in ADHD.

Regarding motor behavior, unobtrusive computer vision effectively captured ADHD-related restlessness that typically requires specialized sensors [82,83]. Accounting for age-related decreases, widespread head, upper body, and facial movement specifically marked the ADHD-Hyperactive status. Neither ADHD-Inattentive nor ASD diagnoses affected the global displacement metrics we used here. Because motor challenges are core to ASD [47], the absence of an ASD-associated signal in these global kinematic measures suggests that they may not capture the more specific, repetitive movement patterns often described in autism [46], which may require a more granular movement classification [46, 84–89]. However, no differences in facial expressivity during naturalistic interactions in ASD are consistent with other recent studies [90].

Developmental and demographic factors proved essential to the behavioral phenotype, with age robustly driving changes across motor, linguistic, and semantic domains alongside sex-specific profiles. Digital biomarkers must rigorously disentangle normative trajectories from pathology. Although cognitive ability varies within ASD, [52, 53], influences language [37, 91], and remains controversial as a pure covariate [92], our primary diagnostic associations remained robust whether Full-Scale IQ was included or excluded (see Fig. S4).

While our automated features correlate with clinical instruments (Fig. S5) and they can offer enhanced presentation-specificity, they are not intended to replace gold-standard assessments. Traditional tools align more closely with categorical DSM-5 definitions (Fig. S6). Instead, automated multimodal analysis provides scalability and objectivity, avoiding inter-rater reliability issues [26]. However, these metrics currently lack qualitative nuance. For instance, current computer vision models quantify displacement but cannot distinguish between a situational fidget, a neurological tic, and an autistic repetitive behavior or mannerism, which raters are trained to reliably differentiate, for example, in the Autism Diagnostic Observation Schedule (ADOS) [93]. Similarly, semantic embeddings capture statistical atypicality but not the underlying pragmatic intent accessible to human raters. Thus, these tools serve as complementary phenotyping instruments, offering a dimensional overlay for subtle, continuous behavioral differences and allowing characterization at scale.

Given the strong developmental focus of this work, it is important to remember that the study remains crosssectional. The observed age-related differences largely align with known developmental milestones for language [94] and movement [95]. However, longitudinal studies are needed to confirm if these automatic behavioral metrics track individual trajectories over time. Automatic methods have already shown promise in predicting future prosody, language, movement, and developmental delay outcomes in ASD [44, 45, 96–98] and ADHD [98, 99].

Additionally, because individual-level records of ADOS or ADI-R administration were unavailable for this specific dataset, we relied on continuous screening instruments (e.g., ASSQ, SRS) rather than gold-standard observational subscores to characterize autism trait severity.

Furthermore, the generalizability of our findings to the broader autism spectrum must be considered with caution. Because the semi-structured interview paradigm inherently depends on verbal communication and sustained social interaction, our ASD cohort included only verbal individuals with mild-to-moderate support needs (see Fig. S7). Consequently, our conclusions on narrative production and speech prosody do not apply to minimally verbal individuals. This mild-to-moderate clinical profile also likely explains why we did not find pronounced motor stereotypies [100], prompting our focus on global kinematics instead, which are better suited to track generalized restlessness. Additionally, our ASD sample is characterized by a particularly high rate of co-occurring ADHD ( 80%). While this reflects the realities of community-based clinical samples [101, 102], it underscores that our findings are most applicable to this specific, highly prevalent, and verbally fluent segment of the neurodivergent population.

Last, our analytic framework assumed that behavioral presentations of ASD and ADHD add up in individuals with both diagnoses. This allowed us to identify the main effects of each condition, in line with the NIMH Research Domain Criteria that aims to parse out biological or cognitive mechanisms [57, 58]. Prior work on smaller samples suggested that ASD and ADHD co-occurrence may interact to exacerbate effects on verbal fluency [103], and cognitive and behavioral traits [104]. The lack of significant interaction effects in the present study provided empirical support that the co-occurring presentation is adequately captured by the sum of the independent main effects, at least for the behavioral metrics analyzed here. Future work on ADHD and ASD should validate these brief, video-based measures longitudinally using wearable sensors and apply them within standardized assessments like the ADOS [93]. More broadly, this work illustrates how brief, standardized clinical interactions can be transformed into multidimensional behavioral measures at scale. A similar approach may be relevant beyond developmental disorders, to include neurological and psychiatric conditions where patients present with altered movement, posture, speech, or language.

## 4. Methods

### 4.1. Clinical Population

This study used video-recorded conversations with 2,529 children (aged 5-22, *M* = 10.08, *SD* = 3.38 years) as part of the Healthy Brain Network (HBN) [30]. The cohort included 1,627 participants with clinician-confirmed ADHD: 722 with inattentive presentation, 107 with hyperactive-impulsive presentation, and 702 with combined presentation. An additional 96 showed ADHD traits without meeting diagnostic criteria for a specific ADHD presentation (DSM-5 ADHD-other/unspecified). The sample also comprised 306 individuals with mild to moderate ASD (mostly level 1 according to DSM-5). Of the 1627 participants with ADHD, 246 (15.12%) had a co-occurring ASD diagnosis, a rate similar to those reported in previous community-based cohorts [101, 102]. The cohort included 211 Typically Developing individuals and 631 with other clinical diagnoses, mostly anxiety and learning disorders. This group did not exhibit different hyperactive, or inattentive traits compared to the Typically Developing controls and only minor difference in autistic traits (Table 1). Furthermore, despite the wide developmental range of the study, differences in age distributions between diagnostic and Typically Developing groups were small or non-significant (Table 1 and Fig. S1). The overall male-to-female ratio was 1.81:1 (1630/899). Detailed demographic and clinical characteristics per diagnosis are available in Table 1.

Of the *N* = 2, 783 participants who enrolled for the interview at the Child Mind Institute and consented to share their data for research purposes, we excluded cases with an incomplete clinical evaluation (*N* = 157) and participants for whom clinicians recommended additional testing to rule out ASD (*N* = 97). The remaining *N* = 2, 529 had a final clinician-confirmed diagnostic consensus and were included in this study. Notably, no participants were excluded entirely due to general audio or video data quality. Instead, if the available interview recording did not allow for the quantification of a specific measure (e.g., a missing answer to a specific question), that measure was assigned a null value for that participant and was excluded only from the specific statistical models evaluating that variable.

Of the 2,529 participants, 1084 (42.86%) were White/Caucasian, 291 (11.47%) Black/African American, 211 Hispanic (8.34%), 50 (1.98%) Asian, and 28 (1.11%) identified as other races. 353 (13.96%) identified as two or more races. 512 (20.25%) participants did not specify, or their race was unknown.

### 4.2. Behavioral Task, Data Acquisition

Naturalistic behavior was recorded during a semi-structured clinical interview following the viewing of an emotive animated short film, “The Present” (https://www.youtube.com/watch?v=3XA0bB79oGc). The film depicts a boy who initially rejects a three-legged puppy given to him by his mom while playing a videogame; in the final scene, it is revealed that the boy himself has a missing leg. The narrative therefore invites viewers to infer the characters’ emotional states and the similarity between them. The subsequent interview was structured to elicit responses related to narrative recall, emotional description, and perspective-taking (Theory of Mind) (see Table S1 for a list of the questions and Table S3 for an overview of related measures). High-definition video (Canon XC15) of the participant’s face and upper body and high-fidelity audio (Røde NT1) were captured simultaneously for automated analysis. Examples for the interview settings and camera view are available at Fig. S8.

### 4.3. Diagnostic Procedure

The HBN is a large-scale biobank that utilizes a community-referred recruitment model to capture the wide heterogeneity inherent in developmental psychopathology. As such, the HBN sample is enriched for individuals meeting DSM-5 criteria.

Diagnoses within the HBN are determined by licensed clinicians at the Child Mind Institute based on a consensus model that integrates multiple sources of information. The primary diagnostic instrument is the semi-structured Kiddie Schedule for Affective Disorders and Schizophrenia (K-SADS) [105] for DSM-5, which is administered to both participants and their parents. The final clinician consensus diagnosis also incorporates behavioral observations during extensive testing, parental and self-reported clinical questionnaires, and the participant’s developmental, educational, and diagnostic history. For ASD, the primary evaluation relies on the KSADS autism module and questionnaires (e.g., SRS-2, SCQ, ASSQ), as well as developmental and prior diagnostic history. We excluded cases where clinicians recommended additional testing to rule out ASD. A limited subset of participants were confirmed with the ADOS (*N* ∼ 20). Further details on the diagnostic process and clinician-administered assessments are available on the original HBN manuscript [30].

For our analysis, clinician-confirmed diagnoses were modeled to assess the impact of specific conditions. ADHD diagnoses were categorized as inattentive and/or hyperactive/impulsive diagnostic status. Participants with a clinician-confirmed diagnosis of ADHD-Combined presentation were considered positive for both hyperactive and inattentive. ASD diagnoses were modeled as a single binary variable (1 for positive, 0 for negative). The vast majority of these individuals (*>* 95%) had mild to moderate presentations (only a few requiring significant support, DSM-5 level 1), consistent with the HBN’s inclusion criteria, which require participants to be verbal (see Fig. S7 for distribution of autism instrument scores). The control group, designated as the Typically Developing Population, consisted of individuals with a confirmed negative diagnostic clinician decision in all DSM-5 categories.

### 4.4. Clinical Instruments

Participants in the HBN are deeply phenotyped using a comprehensive battery of clinical instruments. For this study, we defined ADHD traits using the Strengths and Weaknesses of ADHD Symptoms and Normal Behavior Scale (SWAN) [106], and ASD traits using the Autism Spectrum Screening Questionnaire (ASSQ) [107]. These scales were selected as they showed the strongest unique association with their respective clinician-confirmed diagnoses compared to other available scales (see Fig. S6). Other clinical instruments used in this study include the Mood and Feelings Questionnaire (MFQ, depressive traits) [108], Screen for Child Anxiety Related Disorders (SCARED-P, anxiety traits) [109], and the Child Behavior Checklist (CBCL) [110], with two factors: internalizing and externalizing traits. A full description and statistical analysis of age, sex, IQ, and clinical instrument scores for each diagnostic group is provided in Tables 1 and S2.

### 4.5. IQ score

Cognitive ability was assessed using a composite IQ score. Full-Scale IQ scores were primarily measured using the Wechsler Intelligence Scale for Children (WISC-V) [59] (*N* = 2, 152, mean *±*SD = 100.4 *±* 16.4). Exceptions included early participants who were administered the Wechsler Abbreviated Scale of Intelligence (WASI) [60] (*N* = 30, 98.1 *±* 16.2), and children under age 6 or with known IQ below 70, for whom the Kaufman Brief Intelligence Test (KBIT) [61] was used (*N* = 162, 100.9 *±* 16.8). Similarly, verbal IQ scores were obtained by combining the WISC-VCI (*N* = 2, 152, 104.7 *±* 16.1), WASI-VCI (*N* = 30, 99.1 *±* 15.7), and KBIT-verbal (*N* = 162, 101.3 *±* 16.6) scores. Full-Scale and Verbal IQ Composite scores were available for 2,344 out of the 2,529 participants (92.7%). The main analysis and results were computed using Full-Scale IQ as a covariate. However, the main associations between diagnoses and measured behavior were independent of including IQ as a covariate or not (Fig. S4). Similarly, the results did not change significantly if modeling with Verbal IQ scores instead.

### 4.6. Transcription, Diarization, & Question-Answer extraction

Audio recordings were transcribed using the WhisperX model (v.3.3.0, Whisper model large-v2) [111], generating a raw transcript with word-level timestamps. Validation against a manually transcribed and diarized groundtruth dataset (*N* = 50, author AS) demonstrated a Word Error Rate (WER) of 3.75% *±* 4.59% (M *±* SD). To separate interviewer and participant speech, we utilized a custom prompt with Google’s large language model (gemini-2.0-flash). This context-aware approach yielded a Diarization Error Rate (DER) of 3.16% *±* 3.97%, significantly outperforming standard acoustic-based methods (pyannote/speaker-diarization-3.1 [112] DER: 16.35% *±* 8.78%). The LLM leveraged the semi-structured interview context to infer speaker turns, avoiding acoustic ambiguities caused by pitch and/or speech overlap between children and interviewers. A secondary LLM prompt subsequently isolated specific interview questions and the participant’s corresponding answers. Complete prompts are provided in the Supplementary Sections S2 and S3. The number of valid answers per question is available in Fig. S9.

### 4.7. Description of Behavioral Variables

Various open-source tools were used to obtain objective measures of language, speech prosody, semantic content, and movement from the interview data. An in-depth description of the measures and their extraction methods is provided in Table S3; the distribution of the measures is available in Figure S10. The code for the extraction of measures is available under the link in the Code Availability section.

#### Language and Speech Prosody Measures

We extracted a set of acoustic and linguistic features using the Openwillis toolkit (v.3.0.5) [63]. Openwillis is a Python wrapper developed by Brooklyn Health that automates feature extraction by integrating previously established open-source speech and natural language processing tools for the extraction of objective language and speech measures. See Table S3 for a complete description of all measures and their extraction methods.

#### Age-Normed Semantic Measures

To characterize the semantic content of participants’ answers, we utilized Google’s ‘Gecko’ text embedding model (text-embedding-004) to generate 768-dimensional numerical representations of each response that capture meaning in a high-dimensional semantic space [113]. To distinguish diagnosis-related divergence from typical developmental immaturity, we adopted an age-normed approach. First, we stratified the Typically Developing control sample into three developmentally distinct age bins: Middle Childhood (5-10 years), Early Adolescence (10-14 years), and Late Adolescence/Young Adulthood (14-22 years). Subsequently, for each of the 23 interview questions, we calculated a bin-specific semantic baseline by computing the median embedding of answers provided by TD participants within that specific age range. We then computed the cosine similarity between each participant’s response and the baseline of their corresponding age bin. This yielded a score representing how semantically ‘typical’ the response was. Participants with missing answers were assigned a missing score for that question. The interpretation of this score depends on the question’s content; for narrative-recount questions, higher similarity reflects greater “typical narrative detail,” whereas for questions about emotion, lower similarity may indicate more nuanced or idiosyncratic descriptions. Finally, these 23 individual typicality scores were averaged into conceptually meaningful domains based on the function of the questions. These domains included narrative detail, factual memory, thematic understanding, and perspective taking (e.g., describing one’s own or a character’s emotions, or the similarity between the dog and the kid, and how this relates to the kid’s behavior). For example, the ‘self-emotion description’ score was the average semantic similarity score across answers to questions 14, 17, 20, and 23 (see Table S1 for details on these questions and examples of the ‘Typical Answers’ for each age bin). The Fisher-Z transform (inverse hyperbolic tangent, artanh) was applied to the semantic (cosine) similarity measures prior to regression analysis to approximate normal distributions.

#### Movement measures

We extracted anatomical and facial landmarks from video recordings at the native 30Hz sampling rate of the video recordings using Google’s Mediapipe Holistic model (v.0.10.11) on a local machine. The raw 3D landmark coordinates were filtered to remove low-confidence data based on two criteria applied over a sliding window. First, segments with more than 10% missing values in a 2-second window were excluded. Second, segments exhibiting excessive jitter, defined as a standard deviation of the x-coordinate exceeding empirically-defined thresholds (0.1 for face; 0.01 for pose and landmarks) in 10-frame windows, were also excluded. A landmark point was considered valid for analysis only if it passed both of these checks. To ensure comparability across participants, the filtered data were normalized. Facial landmarks were aligned to a canonical model using an affine transformation [114] to correct for head rotation, translation, and scale. Body landmarks were normalized by the frame-by-frame inter-shoulder distance to render movements invariant to body size and participant’s distance from the camera. Finally, movement was quantified as the frame-to-frame 3D Euclidean displacement for each landmark. These displacement values were then averaged within predefined anatomical regions (e.g., face, mouth, eyes, and body, see Fig. S8) and across all valid frames to yield a single mean movement score per region for each participant. Log of the average movement was used to model the effects of diagnosis on such measures.

### 4.8. Analytical Approach & Statistical Modeling

To test our primary research questions, we modeled the relationship between the measurements of interest and clinical diagnoses using multivariate linear regression. The model isolated the unique contribution of ASD and ADHD diagnostic status while controlling for key developmental and cognitive confounders. For each behavioral metric, the statistical model took the following form:

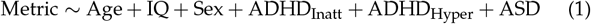

with one model for each dependent variable (e.g., speech rate, lexical diversity), and the predictors were age (in years), a composite IQ score (Full-Scale), Sex (coded binary female=1, male=0), and binary variables for clinician-confirmed diagnoses of ASD, ADHD-Inattentive, and ADHD-Hyperactive (ADHD-Combined presentation was modeled as positive in both).

Every participant counts as *n* = 1, with a set of numbers describing them: Age, IQ, ASD, ADHD-Hyper, ADHD-Inatt. For instance, an 8-year-old boy with an IQ of 90 who presents with ASD and ADHD-Combined is coded as: 8, 90, 0, 1, 1, 1. Models were fitted using the bisquare ‘robust’ linear fitting option in MATLAB (2024b). Variance Inflation Factors (VIF) confirmed that multicollinearity was not a confounding factor (all VIFs *<* 1.3). To quantify and compare the effect of each predictor, we estimated its effect size from the *t*-statistic as follows:

For continuous variables:

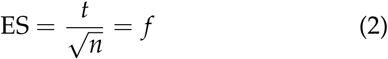

For binary variables:

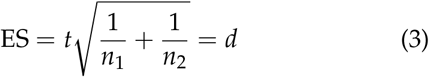

Here *n* is the degrees of freedom, and *n*_1_ and *n*_2_ are the number of data points for each binary group. These effect size measures allow for comparison across different variable types, even when the number of observations varies due to missing data for a specific behavioral metric. For the binary variables (Sex and Disorders), equation (3) is the traditional Cohen’s *d*, and for continuous variables (Age and IQ), equation (2) captures the traditional Cohen’s *f* . In both instances, it is the effect on the outcome variable over the residual unexplained error. Statistical tables for each model of the behavioral outcome measures, including all beta coefficients, standard errors, *t*-statistics, effect sizes, and p-values, are provided in Supplementary Sections S1.2-S1.4. All multivariate model p-values reported in the main manuscript text and figures are corrected for multiple comparisons using Bonferroni correction by multiplying model p-values by the number of tested behavioral features.

To assess diagnostic co-occurrence, we constructed three separate logistic regression models. For each model, one clinical diagnosis served as the binary outcome variable, while the remaining diagnoses were included as predictors, alongside age and sex. Crucially, the specific diagnosis used as the outcome was excluded from the predictor set for that model. We report odds ratios (OR) as the exponents of the beta coefficients (Fig. 1f, Supplement Section S1.1):

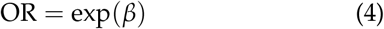

### 4.9. Alternative analysis approaches

To further validate our methodological decisions, we conducted several additional post-hoc sensitivity analyses:

First, although our models accounted for the significant overlap between ADHD and ASD, other co-occurring conditions could have influenced the results. However, we ruled out the impact of child anxiety on our behavioral measures by repeating all analyses while modeling for co-occurring positive anxiety diagnoses, and the associations with ADHD or ASD remained consistent (Fig. S11).

Second, although our reference group included individuals with other non-target DSM-5 diagnoses (*N* = 631), excluding these participants to allow for a direct comparison with Typically Developing youths did not alter the significant associations found for ADHD or ASD.

Third, to mitigate potential bias from the skewed age distribution, we repeated the analysis, limiting it to the 5–14-year-old subset. All results remained the same, except that pitch variation was no longer associated with ADHD-Hyperactive status. To clearly distinguish the variance related to Hyperactive versus Inattentive status, we did not separately code for a Combined presentation, unlike some prior studies [115–117]. When we repeated the analysis by coding with the Combined presentation instead of Hyperactive status, results on movement remained the same, but the effect of Hyperactivity on speech rate and loudness was not there (Fig. S12).

Fourth, our planned analysis treated ASD, ADHD-Hyperactive, and ADHD-Inattentive as independent additive factors (Eq. 1). It is possible that the two conditions interact to produce effects on our outcome measures that are not just their linear superposition. To test for this, we conducted a post hoc follow-up analysis that added interaction terms ASD *×* ADHD-Inatt and ASD *×* ADHD-Hyper to model (1) similar to [103]. We found that neither was statistically significant for any of the outcome measures (all *p*-values ≥ 0.17, after Bonferroni-correction). Additionally, we tried modeling the data using mutually exclusive groups (control, ADHD-only, ASD-only, and ADHD+ASD), as is often done in the literature [54–56]. This approach quantifies differences between groups relative to the control group, but does not assign effects to ASD and ADHD dimensions shared in the combined group. As a result, the distinct separation of effects observed in our primary additive model was lost (Fig. S13), supporting our approach of treating these as independent, overlapping dimensions.

Fifth, to ensure these semantic differences in the ASD group were not merely artifacts of underlying structural language or vocabulary deficits, we conducted an additional post-hoc analysis controlling for clinical language scores (CELF-Screener [77]). The ASD-associated divergence in semantic typicality remained robust (Fig. S3).

Finally, the linear model (1) does not include interaction terms. For age in particular, this means that the effects of the disorders are modeled as a constant gap between a positive and a negative diagnosis. However, for some behavioral features, the gap appears to increase with age (e.g., Fig. 3d, 4b). Such an overall gap can be detected by model (1) even if it increases with age. In a post-hoc analysis, we included the interaction of age with all diagnostic variables. None of the interactions were statistically significant after multiple comparison correction, while the Bayesian Information Criterion increased over the reduced model for most of the measures, suggesting that we are lacking statistical power to warrant the increase in model complexity. Narrowing it to variables with a main age and diagnosis effect suggests that there may be an increasing gap with age for narrative detail in ASD (Fig. 3d, *p* = 0.004, uncorrected). Conversely, differences appear to diminish with age for pitch in ASD (*p* = 0.0009, uncorrected). However, note that these are uncorrected and post-hoc analyses.

### 4.10. Visualization of Distinctive Terms

Semantic embedding analysis of the answers is quantitative and applies to all questions asked. To gain a sense of what it captures, we conducted post-hoc analyses on the significant ASD-question associations. To identify and visualize terms that distinguish ASD from non-ASD participant answers, we employed the scattertext library [62]. The horizontal axis represents the log frequency of each term in the corpus, while the vertical axis represents the term’s association with the ASD group, calculated using a Scaled F-Score (*β* = 2) and normalized via the normal cumulative distribution function to a range of [ −1, 1], as done previously for the comparison of suicidal vs non-suicidal terms [118]. This scoring metric was selected to prioritize recall, where values approaching +1 indicate broad usage within the ASD group, and values approaching *−*1 indicate broad usage within the non-ASD control group. See Fig. S2 for the visualization of answers with a significant ASD association.

## 5. Ethics Statement

The data used in this study were collected between 2016 and 2023. The HBN protocol was approved by the Chesapeake Institutional Review Board. Written informed consent was obtained from participants aged 18 or older, and from legal guardians for participants younger than 18, which included consent for their data to be shared with external researchers for secondary analysis. Access to the data was governed by a Data Transfer and Use Agreement (dated 6/27/2024) between the Child Mind Institute and The City University of New York, while data analysis was approved by the Institutional Review Board of the City University of New York under protocol number 2024-0867-CCNY (dated 12/14/2024).

To protect sensitive Protected Health Information (PHI) inherent in the raw audio and video recordings (i.e., participant faces and voices), all automated AI processing was conducted entirely locally. Feature extraction using computer vision (Google MediaPipe) and speech transcription (WhisperX) was performed on a secure, local machine. No identifiable media or raw biometric data was uploaded to external servers or cloud-based AI services.

## 6. Data Availability

Phenotypical and diagnostic data are available upon request and with the establishment of a Data Use Agreement with the Child Mind Institute at https://fcon_1000.projects.nitrc.org/indi/cmi_healthy_brain_network/. This study used data from Releases 1-11 (publicly available), and Release 12 (yet unreleased). Interview videos are not currently available due to the privacy of the children.

## 7. Code Availability

The full code pipeline used for speech transcription and diarization, as well as to extract language, vocal prosody, semantic embeddings, and body and face landmarks, is publicly available at https://github.com/asortubay/AI_behavioral_analysis.

## 8. Acknowledgments & Funding

NIH-NIMH P50 MH109429, CA Department of Health, grant to Nathan Kline Institute, Milham (PI). Gemini 3.1 Pro was used to assist in shortening our original text in the Introduction, Results and Discussion.

## Notes

### Competing Interest Statement

The authors have declared no competing interest.

### Summary of Updates

Increased sample size to 2,529 from 2,341. Added post-hoc analysis. Updated formatting.

