## Supplementary Materials for "Large-Scale Assessment of Language, Speech, and Movement in Autism and ADHD with AI"

### Supplementary Tables

**Table 1: Interview Questions & Typical Answers**

| No. | Question | 4–10 years | 10–14 years | 14–22 years |
|---|---|---|---|---|
| **Narrative Detail** | | | | |
| 1 | "I hope you enjoyed the last movie. Have you seen it before?" | No. | No? | Yes. |

Continued on next page

**Table 1 – continued from previous page**

| No. | Question | 4–10 years | 10–14 years | 14–22 years |
|---|---|---|---|---|
| 2 | "Can you tell me what happened in the movie? Try to tell the whole story. Remember that stories have a beginning, things that happen, and an ending." | Well, so he was playing a video game. And he was just like, mom. He was playing a video game, and he was sucked into it. And he was like, mm. And then his mom came in the room and, well, came back. from the house and said, and I forgot what she said, but she put a box in front of him and he was like, what is this? Oh my God. And he kept on playing. The mother said, why don't you look what I got for you in the box? And then she got a phone call and the kid was like, for me? And then he opened it and then he saw the puppy and he was very grateful, but then he saw the puppy had no paw and he was like, and he threw him away. And the puppy was just like, and then the puppy went back to him and he just threw him away again. Then the puppy just found like this red ball and the puppy was playing with the ball because it was his first time meeting a ball, I think. And the ball went to the went to the boy and the puppy was like throw the ball for me please and the boy just kicked the ball because he was angry just because yeah and the puppy was playful and kept on running and fell and run fell and run and um The puppy went back to the boy, and the boy, well, no, no, no. And then the boy started smiling, and then he was like, oh, no, no. And then... He kept on smiling, and he was like, okay. And then the puppy went then back to the boy, and the boy got up from his seat, and he didn't have a leg. The same as with the puppy. | So, in the beginning, a boy is playing video games on his couch. And his mom walks over in a box. And he puts the box in front of him, telling him to open it. And at first, he has no interest in it. But then, when he sees what's inside of it, he kind of gets happy. But when he sees that he has a leg missing, he gets all mad, and throws the puppy on the floor. So then, he continued playing video games. And then the dog sees a ball, and he starts chasing the ball around. and kind of limping. So the boy, the dog passed the ball to him, and he kind of kicked it, and the dog chased after it. So after the boy realizes how fun the dog is, he gets up, and but also he has the exact same thing. He lost his leg too. So they go outside and kind of play catch with the dog. | OK. So it started off with the boy. He seemed kind of upset, and he was just playing video games. And it was kind of dark in their house and stuff. And his mom comes home, and she like turns on the lights and everything and she gives him, she gives him the gift in the box so he stops playing and then he opens the box and he like sees a puppy and then he kind of gets excited and then he sees that the puppy doesn't have a leg, like is missing one leg. So then he's like not interested and he kind of just puts it off to the side but the dog doesn't really know what's going on. It's just like excited to be with someone like an owner or something. And then he goes back to playing video games, and the dog is kind of just searching around the house and finds the ball. And you can tell that the dog's kind of having trouble walking and playing with it and stuff. And so then he just kind of watches the owner. He kind of watches as the dog's kind of struggling. but he's playing with the ball and it's kind of making him kind of happy watching him just like play around and he's like still happy. And then he eventually turns off his video game and then it ends when they go outside to play. And you realize that he has some, like he, his leg also is missing. |
| 3 | "Do you remember anything else from the story?" | No. | No. | Yeah, no. Um, no. Okay. |

Continued on next page

**Table 1 – continued from previous page**

| No. | Question | 4–10 years | 10–14 years | 14–22 years |
|---|---|---|---|---|
| **Positive & Negative Preferences** | | | | |
| 4 | "What are some of the things you liked about the movie?" | I liked the dog. I liked the dog. He was really, really cute. | I liked that there was a dog in it and how at the end he was nice to the dog. | I like the message. You can have normal actions and still be impaired in some way and that's kind of motivating and moving. And I like the parallelism between the dog and the boy. And the dog had the same, or at least a really similar disability as the child. But the dog stayed happy throughout. And the kid, he was probably a sad kid, because he only had one leg. But after playing with the dog, he was like, oh, yeah, I can be happy too. I don't have to just stay here and play video games the entire time. |
| 5 | "Is there something you didn't like about the movie?" | That he kicked it. And he didn't like the dog. | There wasn't a whole lot that I didn't like. I mean, like, honestly, I didn't like the fact that he was being mean in the beginning, kicking the dog. | I didn't like the part where he kicked the dog. That was a little bit rude. Other than that, not really. |
| **Factual Memory & Thematic Understanding** | | | | |
| 6 | "Who gave the boy a box?" | His mom. | His mom. | The mom. |
| 7 | "What was in the box?" | A puppy. | A puppy. | A dog. |
| 8 | "What was the boy doing before he got the box?" | Playing video games. | Playing video games. | Playing video games. |
| 9 | "What was the puppy playing with?" | A ball. | A ball. | A ball. |
| 10 | "How are the puppy and the boy the same?" | But they both have one missing leg. | They're both missing a leg. | They both are missing a leg. |
| 11 | "In the movie, who is missing a leg? The boy, the puppy, or no one?" | Both, the boy and the puppy. | Both. | Both. |
| **Emotion Description – Clip 1 (00:53-01:02)** | | | | |
| 12 | "How do you think the puppy was feeling?" | Happy. | Happy, excited. | Happy. |
| 13 | "How do you think the boy was feeling?" | Happy. | Excited, happy. | Happy, excited. |

Continued on next page

**Table 1 – continued from previous page**

| No. | Question | 4–10 years | 10–14 years | 14–22 years |
|---|---|---|---|---|
| 14 | "And how did you feel while you were watching that part?" | Happy. | Happy. | Happy. |
| **Emotion Description – Clip 2 (01:03-01:09)** | | | | |
| 15 | "How do you think the puppy was feeling?" | Sad. | Um, like sad and disappointed. | Kind of probably confused and sad. |
| 16 | "How do you think the boy was feeling?" | Um, mad. | Disgusted and mad. | Angry, upset, disappointed. Yeah. |
| 17 | "And how did you feel while you were watching that part?" | Sad. | I was feeling kind of mad. Yeah. Or just sad. | Um, upset. |
| **Emotion Description – Clip 3 (01:03-01:09)** | | | | |
| 18 | "How do you think the puppy was feeling?" | Sad. | Sad. Like he tried again to like play and all that and then he just kicked him over to get lost. | So I was first kind of excited, like still trying to get attention, and then I'm getting kicked away. |
| 19 | "How do you think the boy was feeling?" | Um, mad. | Um, angry that the dog wouldn't stop annoying him, or annoyed. | Angry. Still mad. |
| 20 | "And how did you feel while you were watching that part?" | Sad. | Kind of mad and a little bit sad. | A little bit sadder because he was actually kind of unleashing his emotions at the puppy. |
| **Emotion Description – Clip 4 (01:14-01:23)** | | | | |
| 21 | "How do you think the puppy was feeling?" | Happy. | Happy. | Happy. |
| 22 | "How do you think the boy was feeling?" | Happy. | Happy. | Happy, too. |
| 23 | "And how did you feel while you were watching that part?" | Happy. | Um... Happy. | Um, happy. |

* Typical Answers are those that have the highest cosine similarity to the median embedding of the age-stratified Typically Developing population.

**Table 2:** **Detailed Statistical Test Results for Table 1**

| Variable | ADHD-Inatt vs TD | ADHD-Hyper vs TD | ASD(ADHD+) vs TD | ASD(ADHD-) vs TD | Other vs TD | ASD(ADHD+) vs ASD(ADHD-) |
|---|---|---|---|---|---|---|
| Age | $Z = 3.31, p = 0.005$ | $Z = -0.90, p = 1.000$ | $Z = 1.99, p = 0.232$ | $Z = 1.62, p = 0.523$ | $Z = 4.39, p < 0.001$ | $Z = -0.54, p = 0.586$ |
| Sex (Distribution) | $\chi^2(1) = 20.76, p < 0.001$ | $\chi^2(1) = 36.45, p < 0.001$ | $\chi^2(1) = 56.59, p < 0.001$ | $\chi^2(1) = 7.43, p = 0.032$ | $\chi^2(1) = 1.62, p = 1.000$ | $\chi^2(1) = 5.34, p = 0.021$ |
| Inattention (SWAN) | $Z = 15.08, p < 0.001$ | $Z = 13.68, p < 0.001$ | $Z = 11.79, p < 0.001$ | $Z = 4.27, p < 0.001$ | $Z = 2.51, p = 0.060$ | $t(261.0) = 3.89, p < 0.001$ |
| Hyperactivity (SWAN) | $Z = 11.49, p < 0.001$ | $Z = 15.75, p < 0.001$ | $Z = 11.83, p < 0.001$ | $Z = 4.01, p < 0.001$ | $Z = -0.53, p = 1.000$ | $t(261.0) = 3.87, p < 0.001$ |
| Autism (ASSQ) | $Z = 10.74, p < 0.001$ | $Z = 11.79, p < 0.001$ | $Z = 16.34, p < 0.001$ | $Z = 10.29, p < 0.001$ | $Z = 3.28, p = 0.005$ | $t(301.0) = -0.09, p = 0.925$ |
| Depression (MFQ) | $Z = 9.55, p < 0.001$ | $Z = 9.41, p < 0.001$ | $Z = 8.59, p < 0.001$ | $Z = 3.29, p = 0.005$ | $Z = 4.59, p < 0.001$ | $Z = 1.94, p = 0.052$ |
| Anxiety (SCARED) | $Z = 6.69, p < 0.001$ | $Z = 5.95, p < 0.001$ | $Z = 7.05, p < 0.001$ | $Z = 5.72, p < 0.001$ | $Z = 6.60, p < 0.001$ | $Z = -1.14, p = 0.253$ |
| Externalizing (CBCL) | $Z = 11.36, p < 0.001$ | $Z = 14.28, p < 0.001$ | $Z = 9.80, p < 0.001$ | $Z = 2.58, p = 0.049$ | $Z = 3.39, p = 0.004$ | $Z = 3.47, p < 0.001$ |
| Internalizing (CBCL) | $Z = 9.83, p < 0.001$ | $Z = 9.46, p < 0.001$ | $Z = 10.85, p < 0.001$ | $Z = 6.11, p < 0.001$ | $Z = 7.45, p < 0.001$ | $Z = 0.40, p = 0.686$ |
| Full-Scale IQ | $t(1524.0) = -6.78, p < 0.001$ | $t(973.0) = -5.22, p < 0.001$ | $t(427.0) = -7.70, p < 0.001$ | $t(251.0) = -4.03, p < 0.001$ | $t(763.0) = -4.97, p < 0.001$ | $t(280.0) = -0.78, p = 0.433$ |
| Verbal IQ | $Z = -5.38, p < 0.001$ | $Z = -4.35, p < 0.001$ | $t(427.0) = -7.19, p < 0.001$ | $t(251.0) = -4.37, p < 0.001$ | $Z = -4.28, p < 0.001$ | $t(280.0) = -0.38, p = 0.702$ |
| Interview Length | $Z = 0.02, p = 1.000$ | $Z = -0.97, p = 1.000$ | $Z = 1.33, p = 0.925$ | $Z = 2.03, p = 0.210$ | $Z = 0.52, p = 1.000$ | $Z = -1.22, p = 0.224$ |

Statistical differences for continuous measures were assessed using independent samples t-tests (for normally distributed data) or Wilcoxon rank-sum tests (for non-normally distributed data), following normality checks via the Kolmogorov-Smirnov test. Categorical comparisons were performed using Chi-square tests. * $p < 0.01$, ** $p < 0.001$ compared to the Typically Developing group (Bonferroni corrected for 5 comparisons). † $p < 0.01$, †† $p < 0.001$ for the comparison between ASD(ADHD+) and ASD(ADHD-).

**Table 3:** **Description of Behavioral Variables**

| Group | Measured Aspect | Variable Name* | Measurement Description | Extraction Method |
|---|---|---|---|---|
| **Language Ability Measures** | | | | |
| Structural Language | Amount of Speech | `speech_length_words` | Total subject speech length in words | Total number of words spoken by the subject in the transcribed & diarized transcript. Log (ln) of total words was used to approximate a normal distribution. |
| Structural Language | Speech Rate | `words_per_min` | Subject speech rate, in words per minute | Total subject words divided by subject speech length in minutes |
| Structural Language | Filler Word Usage | N/A | Number of filler words per minute of subject speech | Total no. of fillers ('um', 'uh', 'er', 'ah', 'like', 'you know', 'so', 'okay', 'right') in subject speech divided by subject speech length in minutes |
| Structural Language | % of unique words | `unique_word_percentage` | Percentage of unique words in total subject speech | 100 - repeated words percentage, measured as the percentage of words in each utterance that are repeated. This is calculated by using a sliding window of 10 words to adjust for utterance length. |

Continued on next page

**Table 3 – continued from previous page**

| Group | Measured Aspect | Variable Name* | Measurement Description | Extraction Method |
|---|---|---|---|---|
| Structural Language | Lexical Diversity | `mattr_10` | Average Moving Average Type Token Ratio with a window size of 10 | Moving Average Type Token Ratio (MATTR), computed using the average of TTRs (computed as t/w, where t is the number of unique terms/vocab, and w is the total number of words) over successive segments of a text by taking the estimate TTR for tokens 1 to n, 2 to n+1, 3 to n+2, and so on until the end of the text (where n is window size), then taking the average. Extracted here using the library `lexicalrichness` with a window size of 10 on the subject utterances. |
| Structural Language | Between Word Coherence | `word_coherence_mean` | Average measure of the semantic similarity (cosine) of each word to the preceding word using LLMs | Average semantic similarity of each word to the immediately preceding word, computed by the cosine similarity between the BERT embeddings (`bert-base-cased`) of each word. |
| Pragmatic Language | Subject Conversation Presence | `speech_percentage` | Time spent speaking by the subject over the total time | Sum of all the subject speaking times divided by the total conversation time. |
| Pragmatic Language | Self-Focused Speech | `first_person_percentage` | Percentage of pronouns in the subject speech that are first-person singular pronouns | Percentage of ("I", "me", "my", "mine", "myself") occurences in the subject speech divided by total pronoun occurences, identified by nltk's PRP (Personal Pronouns) and PRP$ (Possesive Pronouns). |
| Pragmatic Language | Question-Answer Coherence | `turn_to_turn_tangentiality_mean` | Average semantic similarity of the current speaker's turn to the previous turn of the other speaker | Average semantic similarity of each subject turn to the immediately preceding interviewer turn, computed by the cosine similarity between the `sentence-transformers/all-MiniLM-L` text embeddings of each turn. |
| **Semantic Measures** | | | | |

Continued on next page

**Table 3 – continued from previous page**

| Group | Measured Aspect | Variable Name* | Measurement Description | Extraction Method |
|---|---|---|---|---|
| Memory | Factual Memory | Answer Typicality | Answer Typicality when asked about facts in the movie | Semantic similarity (cosine similarity) of each participant answer's 768-dimensional text embedding (Google's `text-embedding-004`), with the corresponding age-matched median 'typical' answer embedding. The Fisher-Z transform (inverse hyperbolic tangent, atanh) was applied to the semantic (cosine) similarity measures prior to regression analysis to approximate normal distributions. |
| Social Judgment & Salience | External Emotion Description | Answer Typicality | Answer Typicality when asked about dog/child emotions in the movie clips | As above. |
| Emotion Description | Self Emotion Description | Answer Typicality | Answer Typicality when asked about self-emotions in the movie clips | As above. |
| Narrative Production, Comprehension & Thematic Understanding | Narrative Detail | Answer Typicality | Answer Typicality when asked to give a detailed recall of the movie | As above. |
| Thematic Understanding | Thematic Understanding | Answer Typicality | Answer Typicality when asked about the similarities between the dog and the kid, and their shared experience | As above. |
| Social Judgment & Salience | Positive Preferences | Answer Typicality | Answer Typicality when asked about what they liked in the movie | As above. |
| Social Judgment & Salience | Negative Preferences | Answer Typicality | Answer Typicality when asked about what they did not like in the movie | As above. |
| **Vocal Prosody Measures** | | | | |
| Prosody | Pitch | `f0_mean` | Vocal Pitch (f0, Hz) | Average and standard deviation for the frame-wise subject fundamental frequency (f0), extracted in the 75-500 Hz range using Parselmouth's praat tool. |
| Prosody | Pitch Variation | `f0_stddev` | Pitch variation (std(f0)) | As above. |
| Prosody | Loudness | `loudness_mean` | Vocal loudness in dB | Average and standard deviation for the frame-wise subject loudness (in dB), extracted using Parselmouth's praat tool. |

Continued on next page

**Table 3 – continued from previous page**

| Group | Measured Aspect | Variable Name* | Measurement Description | Extraction Method |
|---|---|---|---|---|
| Prosody | Loudness Variation | `loudness_stddev` | Loudness variation (std(dB)) | As above. |
| Prosody | Dysphonia | `cpp_mean` | Mean cepstral peak prominence | Average framewise subject Cepstral Peak Prominence (Fraile & Godino-Llorente, 2014). |
| Prosody | Breathiness | `gne_ratio` | Glottal-to-noise excitation ratio | Glottal-to-noise excitation ratio (Godino-Llorente et al, 2010), extracted using Parselmouth's tool. |
| **Movement Measures** | | | | |
| Movement | Facial Movements | N/A | Total movement in the video, per region. | Anatomical and facial landmarks were extracted using Google's Mediapipe Holistic model (model_complexity=2). Then filtered to remove low-confidence or jittery segments and normalized to correct for head orientation and body size. Movement was quantified as the frame-to-frame 3D Euclidean displacement, averaged within specific anatomical regions for each participant. Finally, these regional averages were log-transformed to approximate a normal distribution. |
| Movement | Eye Movements | N/A | Total movement in the video, per region. | As above. |
| Movement | Mouth Movements | N/A | Total movement in the video, per region. | As above. |
| Movement | Upper Body Movements | N/A | Total movement in the video, per region. | As above. |

*Variable Names for the specific measures that were extracted using the OpenWillis tools. For Language Ability Measures, using `speech_characteristics` function, with the optional settings `min_turn_length = 5, min_coherence_turn_length = 5, option = 'coherence'`. For Vocal Prosody Measures, using the `vocal_acoustics` function, with the optional `voiced_segments = True` setting to only compute measures in voice segments longer than 100 ms.

### Supplementary Figures

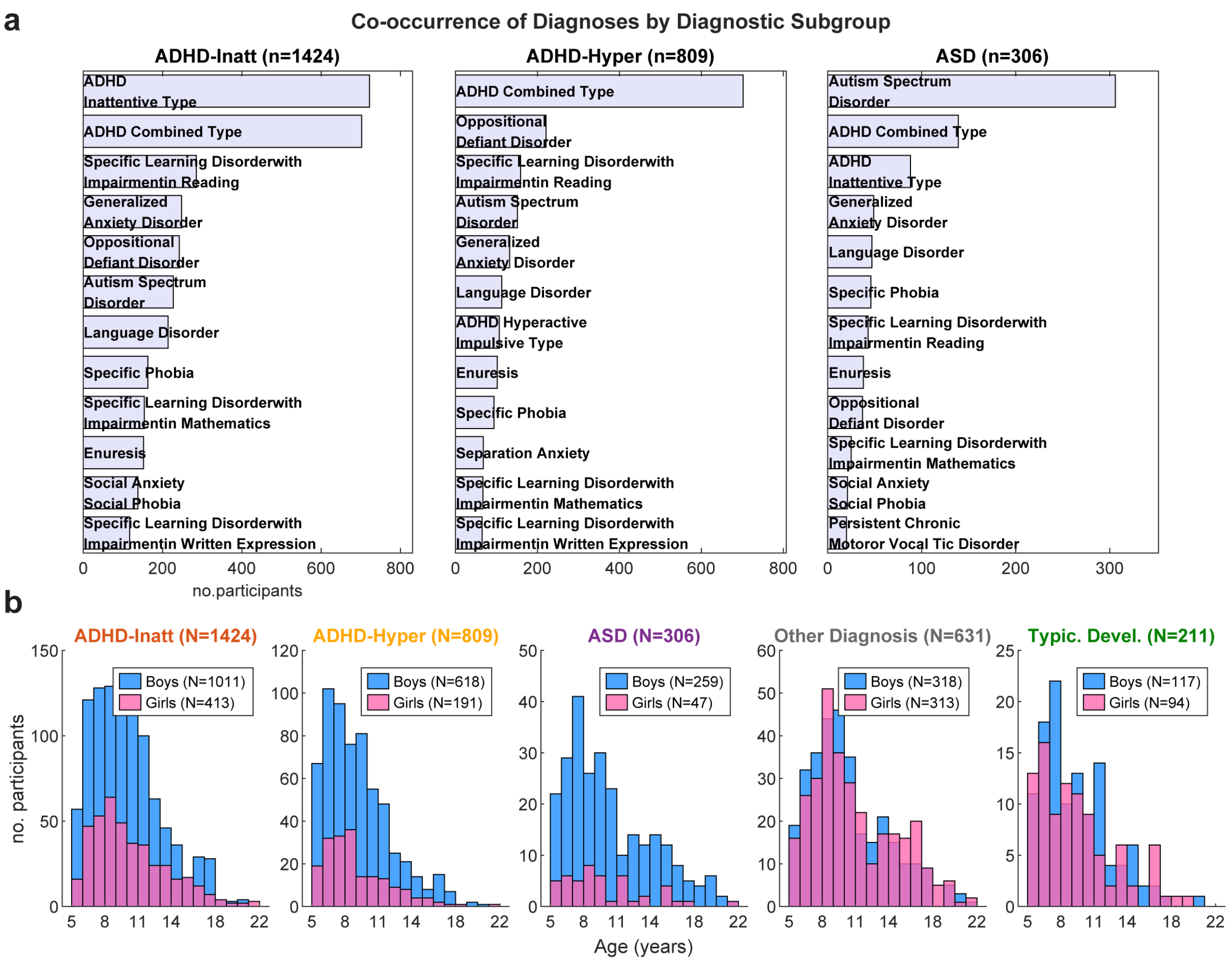


**Figure 1: Co-occurring diagnoses and age distributions.** a) Co-occurring diagnoses (Top 12) in each of the three diagnostic groups of interest. b) Age-Sex distribution across the diagnostic groups. ADHD-Inatt and ADHD-Hyper contain participants with an ADHD-Combined presentation. Other Diagnosis group contains participants with some positive diagnosis, excluding those with ADHD or ASD.

How do the features used here relate to established clinical instruments based on questionnaires? As one may expect, our measures of movement, language, and speech correlate strongly with these established metrics (Figure S5a, b). When using standard instruments, it appears that ADHD-Inattention is also associated with elevated motor activity (Fig. S5c). Thus, our movement measures appear to be more specifically associated with the ADHD-Hyperactive status (cf. Fig. 5c with S5c). On the other hand, when using standard instruments, it appears that ASD has a lesser effect on language use (Fig. S5d). Therefore, our features related to social communication appear to be more sensitive to the ASD presentation (cf. Fig. 3e with S5d).

To select the most specific and robust trait measures for our analyses in Figures 1-2, we evaluated which of these scales best and most uniquely reflected the corresponding clinician-confirmed diagnosis. We ran separate multivariate regression models for each clinical instrument, entering the diagnoses as predictors. This approach allowed us to see the instruments most uniquely associated with each diagnosis. As shown here, the SWAN Inattention and Hyperactivity subscales and the ASSQ emerged as the strongest and most

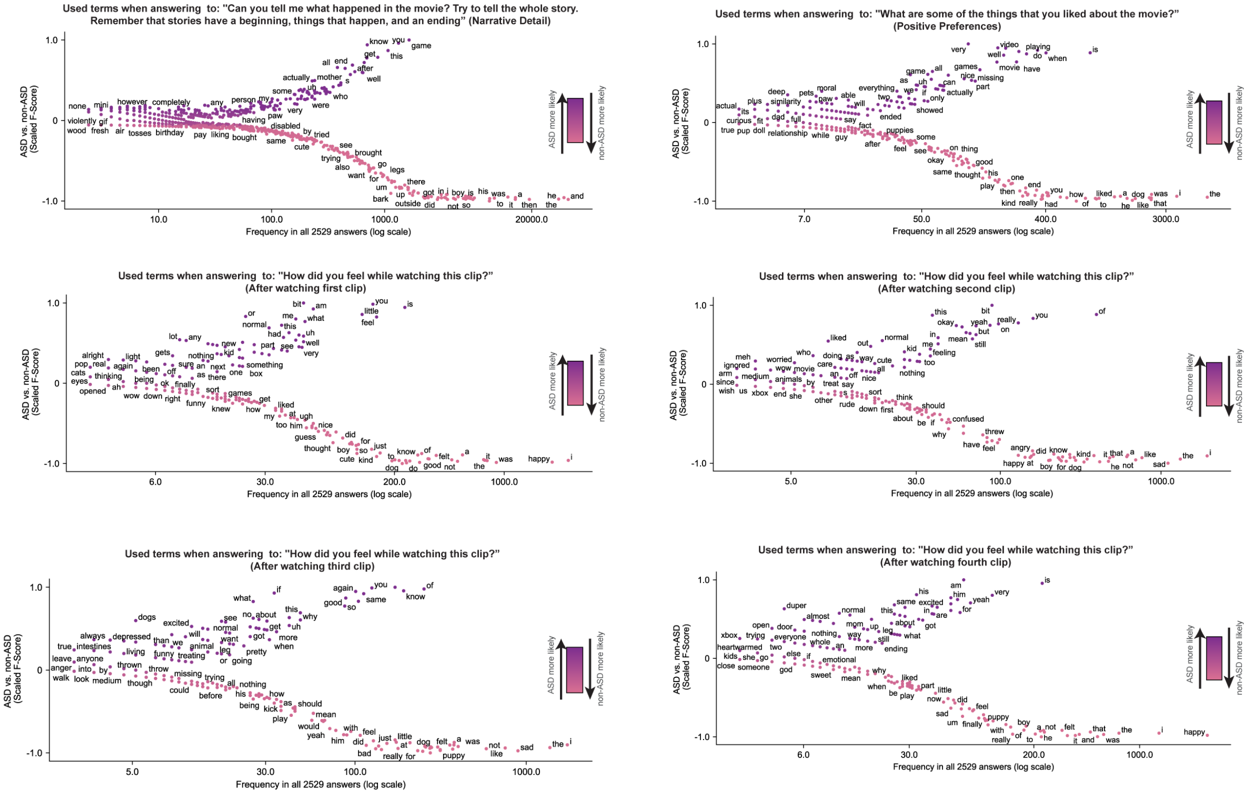


**Figure 2: Scattertext visualization of used terms when answering specific questions.** We only did this post-hoc analysis to add interpretability to the found effects with the semantic analysis; we show here the questions that showed a significant effect of ASD.

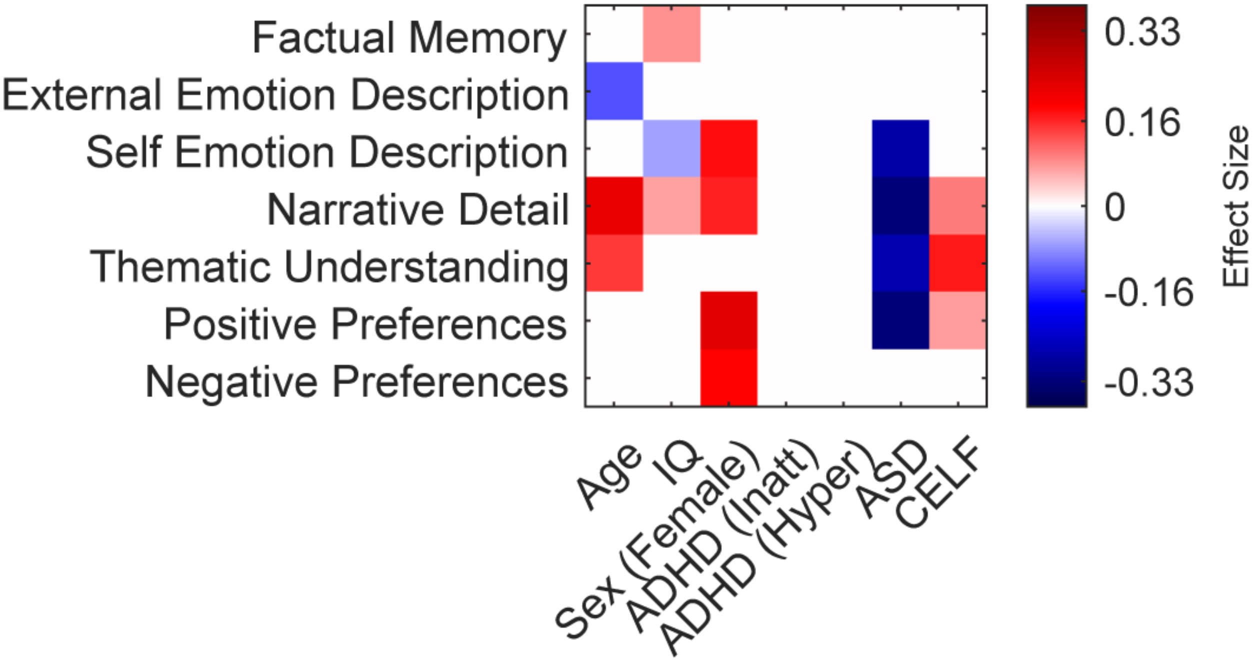


**Figure 3:** **Including clinical language scores as covariate does not remove ASD-semantic associations.** Effect size matrices for semantic outcomes, after including the Clinical Evaluation of Language Fundamentals-Screener Total score as covariate, to rule out possible language deficits that may lead to atypical answers in the ASD population. The main ASD associations remain consistent. Effect sizes are estimated using multivariate regression, with $p < 0.01$ after Bonferroni correction.

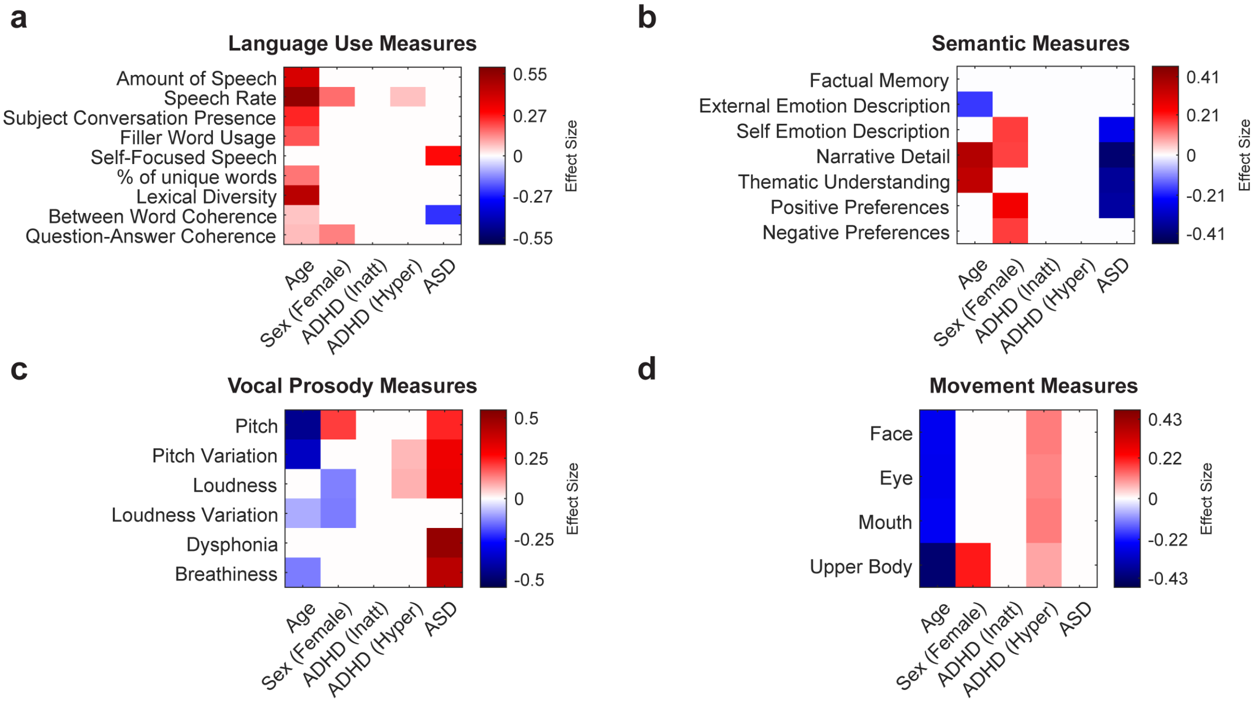


**Figure 4:** **Results are independent of including IQ as a covariate.** Effect size matrices for primary behavioral outcomes after excluding IQ as a covariate. The models (a through d) are identical to the primary multivariate models presented in the main text, but without modelling IQ as a covariate. Effect sizes are estimated using multivariate regression, with $p < 0.01$ after Bonferroni correction.

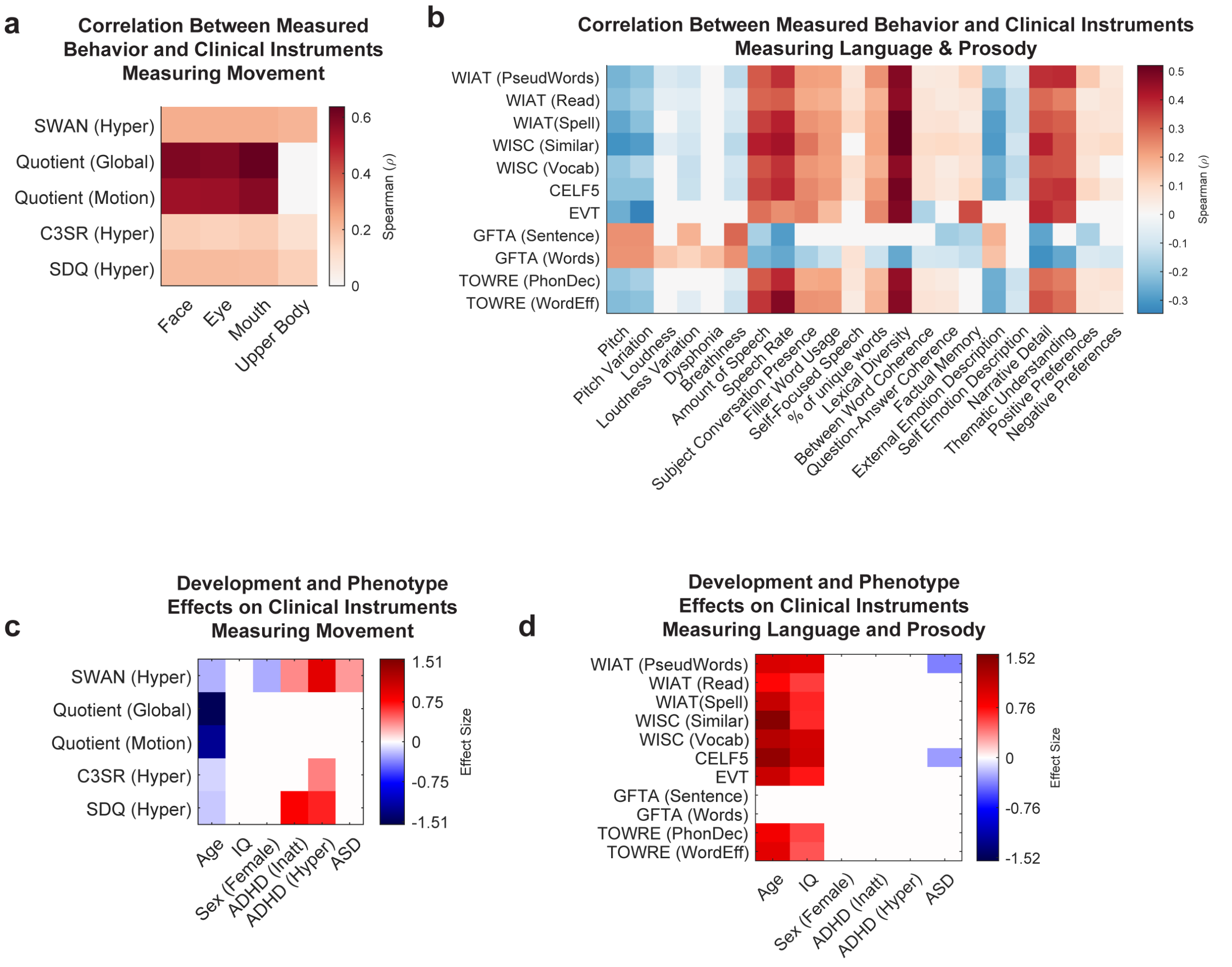


**Figure 5:** **Comparing Clinical Instruments to measured behavior.** Spearman correlation between automatically measured behaviors and the clinical instrument scores measuring movement (a), and language use/prosody (b). c-d) Effects of development and diagnosis on such clinical instrument scores. Effect sizes in c, d measured using multivariate models, significance at $p < 0.01$ after Bonferroni correction, white otherwise. For a and b, significant at a cutoff of $p < 0.05$, uncorrected, white otherwise.

specific associations for their corresponding clinician-confirmed diagnoses. Therefore, we selected these scales for all subsequent analyses where continuous measures of ADHD and ASD traits were required:

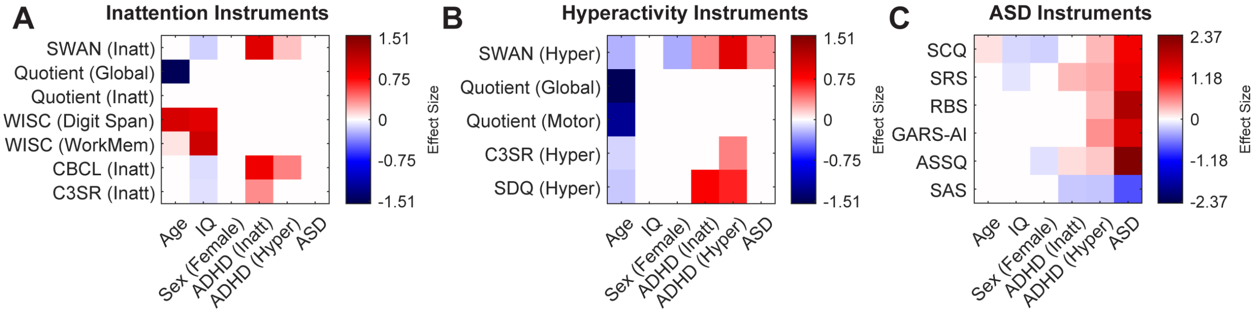


**Figure 6: Unique association of clinical questionnaire scales with clinician-confirmed diagnoses.** The effect size matrices reveal which clinical instruments are more uniquely explained by specific positive diagnoses while accounting for development and demographic effects.

Given the high rates of anxiety in both ADHD and ASD, and the potential for a semi-structured interview with an unfamiliar clinician to elicit anxious behaviors, we modeled this as a key potential confound. Behaviors associated with anxiety, such as motor restlessness, social reticence, or altered vocal tone, could overlap with the objective behavioral markers under investigation and potentially obscure the specific effects of ADHD and ASD. To address this and confirm the robustness of our primary findings, we repeated our primary multivariate regressions, adding a binary variable for any clinician-confirmed DSM-5 anxiety diagnosis as an additional covariate ($N = 909$). This allowed us to test whether the observed associations between our objective behavioral markers and ADHD/ASD diagnoses remained significant after statistically controlling for the influence of anxiety. We find that all the associations between ASD and/or ADHD with our behavioral measures are independent of whether we model anxiety or not:

Throughout the work presented in this manuscript, clinician-diagnosed ADHD-Combined presentations were modelled as positive in both ADHD-Hyperactive and ADHD-Inattentive status. This was done to maximize the number of participants in the modelled group, while maintaining the distinction of the two main inattentive and hyperactive diagnostic statuses. Previous studies on the HBN population have modelled this overlap by considering the ADHD-Combined separate from the ADHD-Inattentive presentations 115–117. We repeated the analyses on the behavioral features by following this approach, and the results were mostly consistent our associations with the ADHD-Hyperactive presentation:

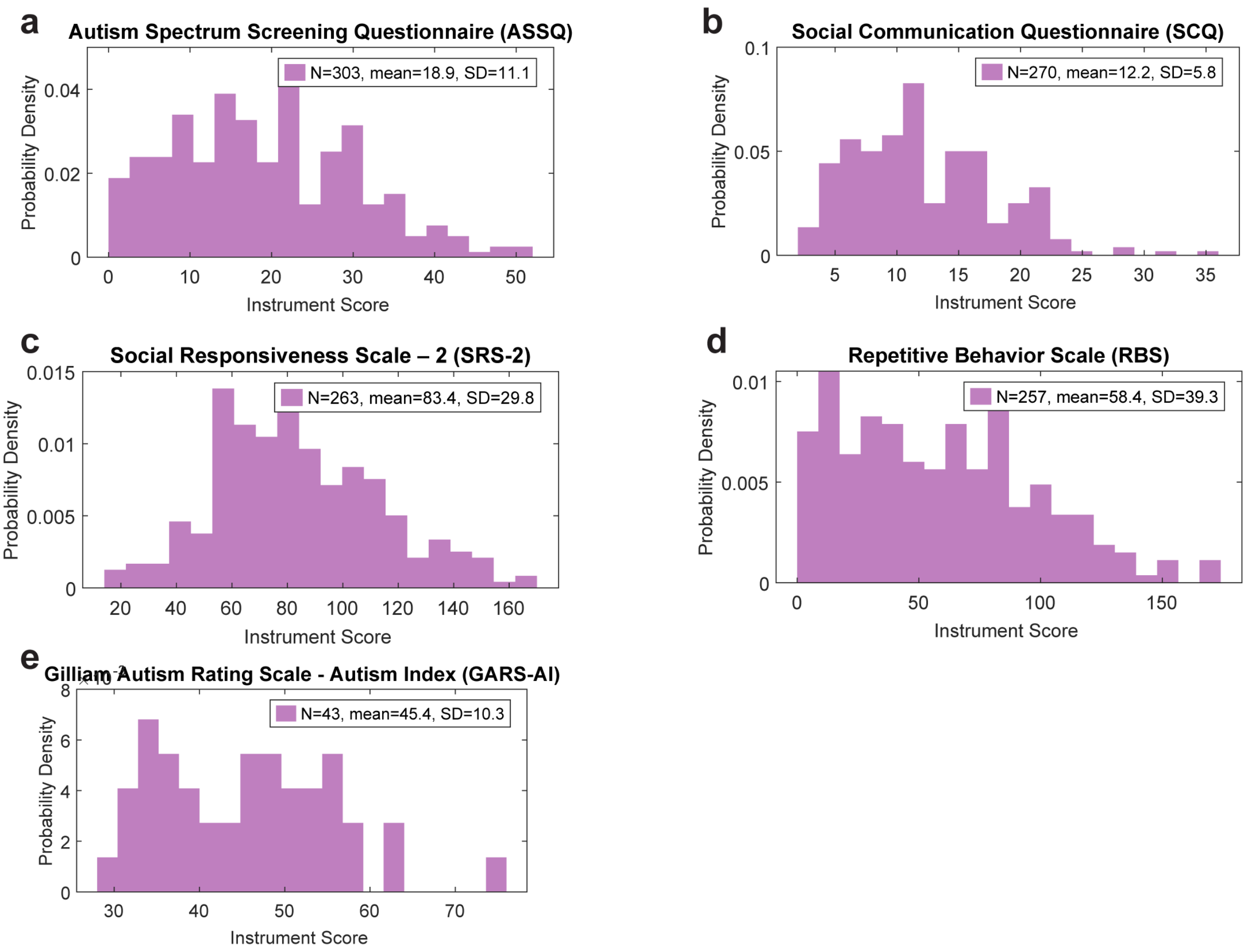


**Figure 7: Distribution of Autism Scores in the ASD-positive population.** The Healthy Brain Network's deep phenotyping protocol provides multiple parent- and self-report questionnaires to measure traits associated with ASD. The ASD population is mainly composed of mild-to-moderate cases.

*[Removed to comply with BiorXiv Policy]*

**Figure 8: Interview video examples, face landmarks, and validity.** The interviews were recorded in different locations and under different lighting conditions. However, all the videos showed the face and upper torso. b) 468 face landmarks tracked by Mediapipe. We characterized total movements for the whole face (blue), only the eyes (magenta), and the mouth (green). c) Most participants showed their face and torso for most part of the video ($N = 2157$ and $N = 2335$ participants were missing 10% or less of the data for face and pose, respectively); however, the amount of available data was less overall for the face landmarks than for the pose landmarks, as participants covered their face or looked away from the camera.

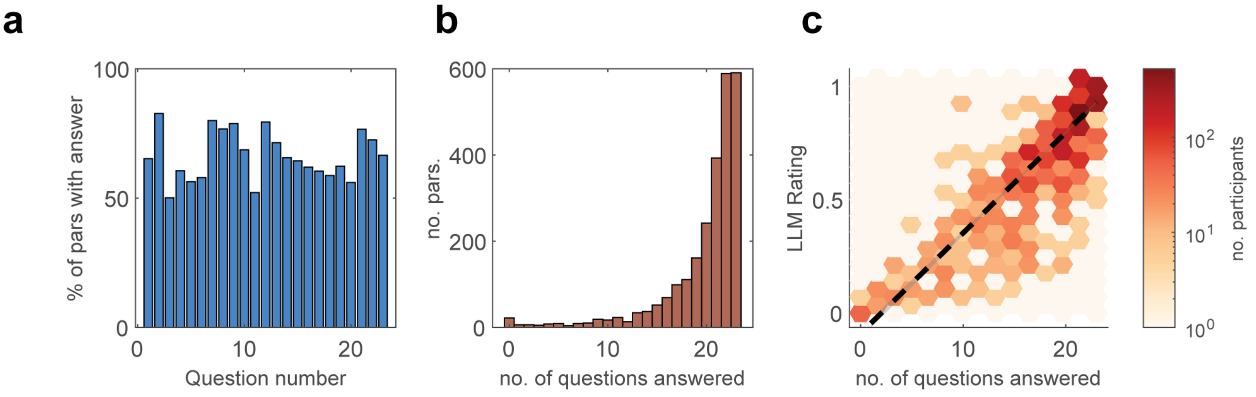


**Figure 9: Validity of interview questions.** a) Not all questions were asked to every participant, or the answer was not available. On average, 66.3% of participants answered each question. b) Each participant answered 19.9 questions on average. C) The number of questions answered by each participant is correlated with the LLM's rating of the conversation completion ($\rho(2527) = 0.80$, $p < 0.001$), see Supplementary Section S3 for rating prompt.

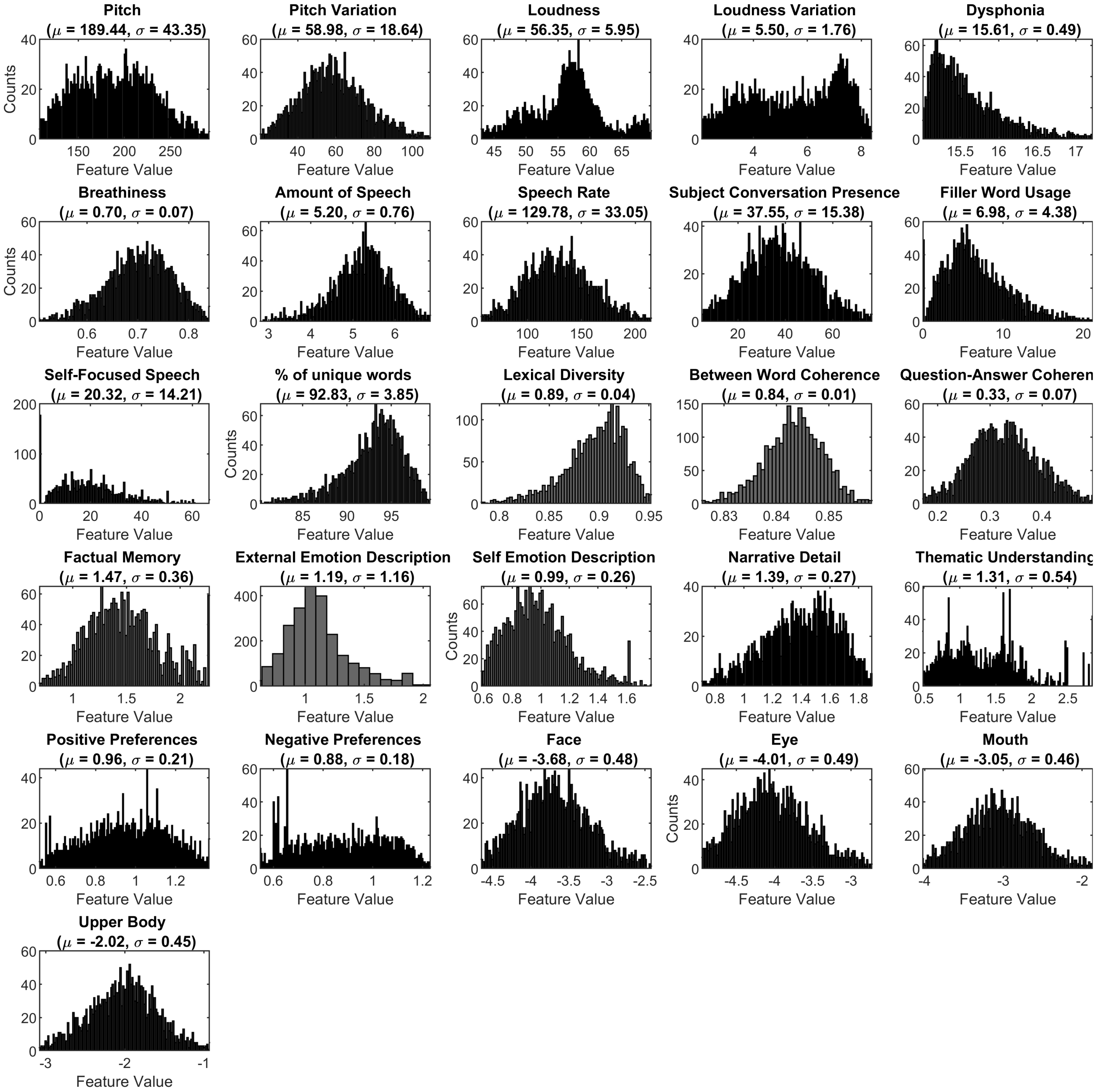


**Figure 10:** **Distribution of Behavioral Variables.** A complete description of variables and their units is available in Table S3.

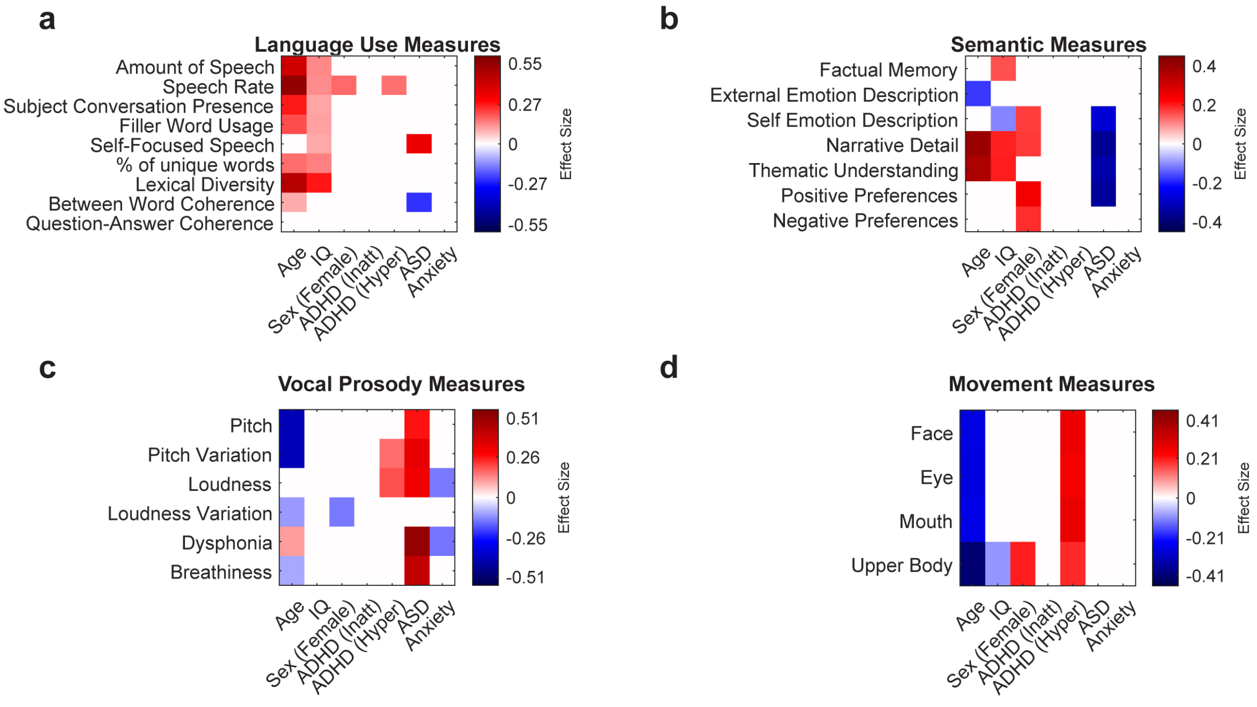


**Figure 11: Results are independent of anxiety diagnoses.** Effect size matrices for primary behavioral outcomes after controlling for co-occurring anxiety. The models (a through d) are identical to the primary multivariate models presented in the main text, with the addition of a binary covariate for any clinician-confirmed anxiety diagnosis ($N = 909$). Effect sizes are estimated using multivariate regression, with $p < 0.01$ after Bonferroni correction.

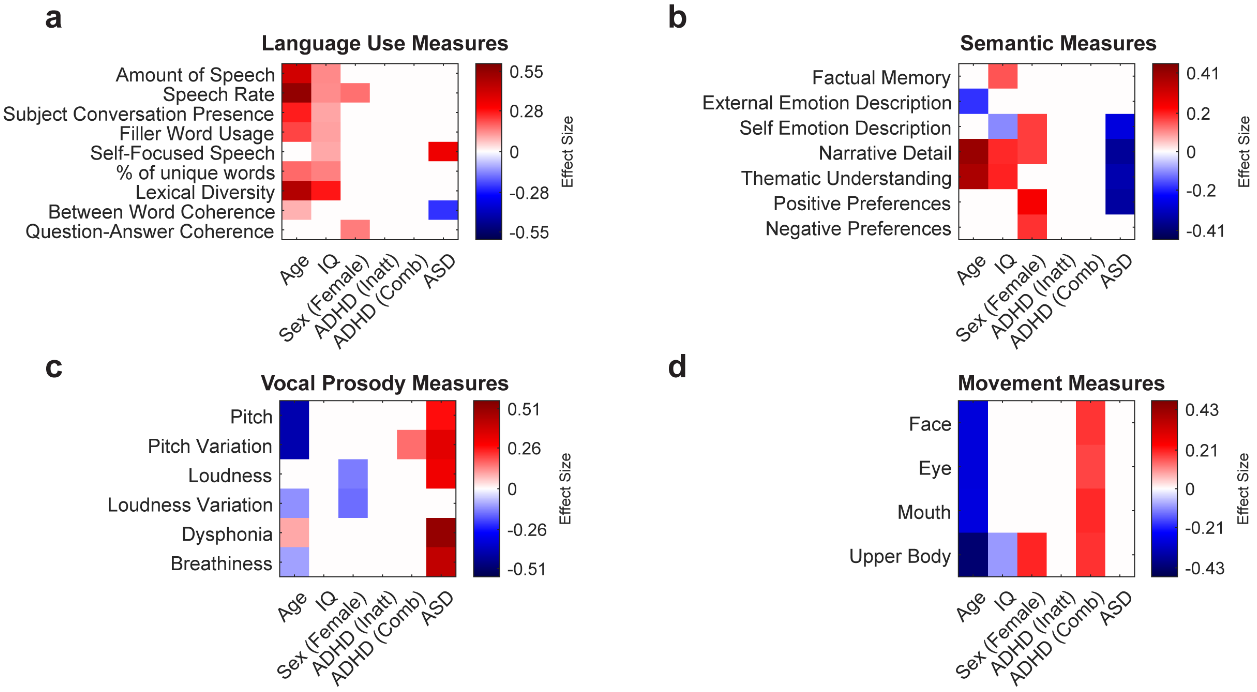


**Figure 12: Results are independent of how the ADHD presentations are coded.** Effect size matrices for primary behavioral outcomes after modelling ADHD-Combined and ADHD-Inattentive presentations separately. The models (a through d) are identical to the primary multivariate models presented in the main text, but ADHD was modelled separately as the ADHD-Combined ($N = 702$) and ADHD-Inattentive ($N = 722$) presentations. Effect sizes are estimated using multivariate regression, with $p < 0.01$ after Bonferroni correction.

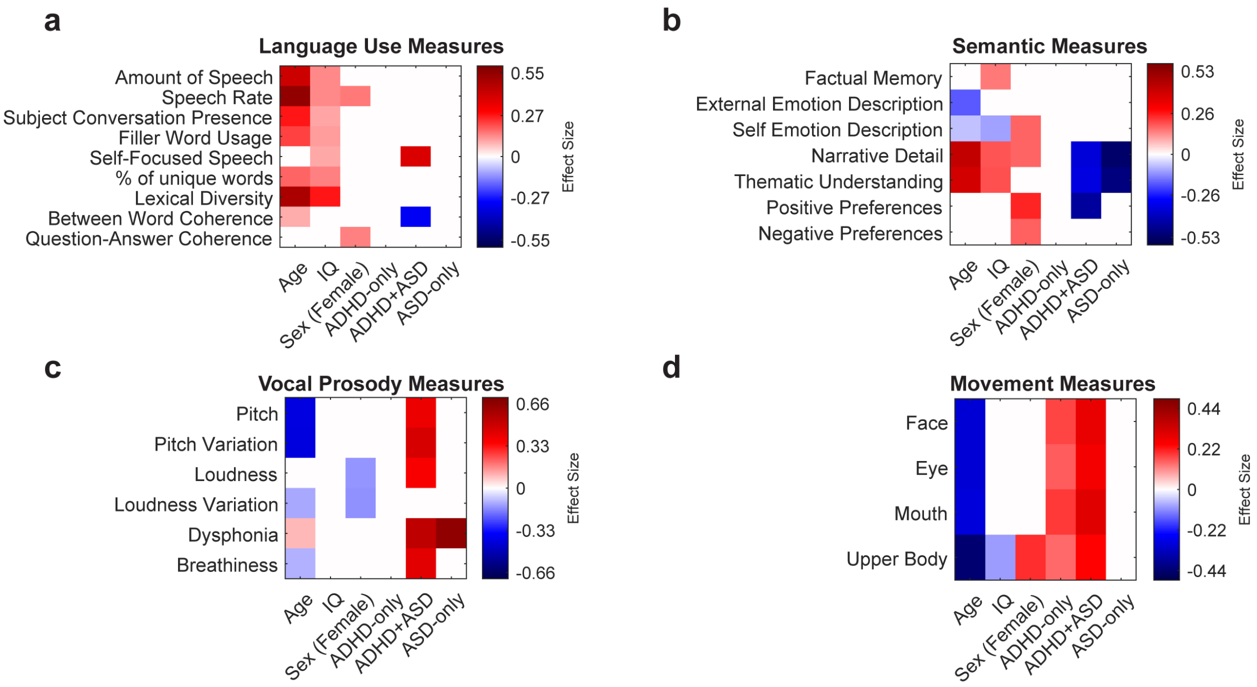


**Figure 13: Modeling ADHD-only, ASD-only, and ADHD+ASD separately does not paint a clear picture.** Effect size matrices for primary behavioral outcomes when treating the diagnosis as a single categorical variable with four mutually exclusive diagnostic outcomes (0:control, 1:ADHD-only, 2: ASD-only, 3:ADHD+ASD). This is equivalent to conventional one-way ANOVA with covariates (ANCOVA). The condition-specific digital phenotype associations observed in the primary model are lost, with effects instead distributed across both the disorder-unique and combined categories. Statistical power is also reduced compared to the primary analysis method in the main text. Effect sizes are estimated using multivariate regression, with $p < 0.01$ after Bonferroni correction.

### Supplementary Section S1. Multivariate Model Reports

$p$-values reported in the tables below appear uncorrected for multiple comparisons unless otherwise specified. In the main manuscript, and in the reported corrected $p$-values below, all $p$-values are corrected for multiple comparisons using Bonferroni correction by multiplying these $p$-values by the number of behavioral features tested. Specifically, in Figures 2-5, Effect Size matrices show significant effect sizes in color with a cutoff of $p < 0.01$ on the corrected $p$-value, and are left blank otherwise. See Methods for the explanation on analytical models.

#### Supplementary Section S1.1. Diagnoses

Detailed model reports for panel f in Figure 1.

**Results for Outcome: ASD**

| Predictor | Odds Ratio (OR) | 95% CI | p-value (corrected) |
|---|---|---|---|
| Age | 1.042 | [1, 1.08] | 0.0887 |
| Sex (Female) | 0.339 | [0.245, 0.471] | 2.95E-10 |
| ADHD (Inatt) | 1.736 | [1.29, 2.34] | 0.000889 |
| ADHD (Hyper) | 1.715 | [1.3, 2.26] | 0.000401 |

**Results for Outcome: ADHD (Inatt)**

| Predictor | Odds Ratio (OR) | 95% CI | p-value |
|---|---|---|---|
| Age | 1.066 | [1.04, 1.09] | 1.53E-06 |
| Sex (Female) | 0.681 | [0.567, 0.818] | 1.53E-06 |
| ADHD (Hyper) | 9.264 | [7.34, 11.7] | 2.04E-78 |
| ASD | 1.734 | [1.29, 2.34] | 0.000314 |

**Results for Outcome: ADHD (Hyper)**

| Predictor | Odds Ratio (OR) | 95% CI | p-value |
|---|---|---|---|
| Age | 0.831 | [0.803, 0.859] | 3.73E-27 |
| Sex (Female) | 0.565 | [0.456, 0.7] | 4.93E-07 |
| ADHD (Inatt) | 9.643 | [7.62, 12.2] | 6.90E-79 |
| ASD | 1.716 | [1.3, 2.27] | 0.000460 |

#### Supplementary Section S1.2. Language Use Measures

Detailed model reports for panels d & e in Figure 2.

**Results for Feature: Amount of Speech**
Model Fit: $R^2 = 0.158$, Adjusted $R^2 = 0.156$ | Overall Model: $F(6, 2288) = 71.440$, $p = 7.98 \times 10^{-82}$

| Predictor | Coefficient ($\beta$) | Std. Error | t-stat | Effect Size | p-value | p-value (corrected) |
|---|---|---|---|---|---|---|
| (Intercept) | 3.76 | 0.103 | 36.6 | NaN | 2.23E-231 | 2.00E-230 |
| Age | 0.0918 | 0.00475 | 19.3 | 0.404 | 4.58E-77 | 4.12E-76 |
| IQ | 0.00541 | 0.000824 | 6.57 | 0.137 | 6.28E-11 | 5.66E-10 |
| Sex (Female) | 0.0522 | 0.0287 | 1.82 | 0.0797 | 0.0686 | 0.617 |
| ADHD (Inatt) | 0.0367 | 0.0301 | 1.22 | 0.0513 | 0.223 | 2.01 |
| ADHD (Hyper) | 0.00475 | 0.0322 | 0.148 | 0.00655 | 0.883 | 7.94 |
| ASD | 0.0122 | 0.0421 | 0.289 | 0.0186 | 0.773 | 6.95 |

**Results for Feature: Speech Rate**
Model Fit: $R^2 = 0.245$, Adjusted $R^2 = 0.243$ | Overall Model: $F(6, 2288) = 123.727$, $p = 9.68 \times 10^{-136}$

| Predictor | Coefficient ($\beta$) | Std. Error | t-stat | Effect Size | p-value | p-value (corrected) |
|---|---|---|---|---|---|---|
| (Intercept) | 48.5 | 4.54 | 10.7 | NaN | 5.04E-26 | 4.54E-25 |
| Age | 5.56 | 0.21 | 26.4 | 0.553 | 1.01E-134 | 9.13E-134 |
| IQ | 0.23 | 0.0364 | 6.31 | 0.132 | 3.36E-10 | 3.02E-09 |
| Sex (Female) | 4.96 | 1.27 | 3.92 | 0.171 | 9.22E-05 | 0.000830 |
| ADHD (Inatt) | -0.361 | 1.33 | -0.271 | -0.0114 | 0.787 | 7.08 |
| ADHD (Hyper) | 5.26 | 1.42 | 3.7 | 0.164 | 0.000224 | 0.00202 |
| ASD | -4.16 | 1.86 | -2.24 | -0.144 | 0.0254 | 0.228 |

**Results for Feature: Subject Conversation Presence**
Model Fit: $R^2 = 0.077$, Adjusted $R^2 = 0.074$ | Overall Model: $F(6, 2288) = 31.598$, $p = 1.1 \times 10^{-36}$

| Predictor | Coefficient ($\beta$) | Std. Error | t-stat | Effect Size | p-value | p-value (corrected) |
|---|---|---|---|---|---|---|
| (Intercept) | 12.7 | 2.42 | 5.26 | NaN | 1.60E-07 | 1.44E-06 |
| Age | 1.42 | 0.112 | 12.7 | 0.266 | 8.24E-36 | 7.41E-35 |
| IQ | 0.0964 | 0.0194 | 4.97 | 0.104 | 7.25E-07 | 6.53E-06 |
| Sex (Female) | 0.636 | 0.675 | 0.942 | 0.0412 | 0.346 | 3.12 |
| ADHD (Inatt) | 1.22 | 0.709 | 1.72 | 0.0724 | 0.086 | 0.774 |
| ADHD (Hyper) | -0.343 | 0.758 | -0.452 | -0.0201 | 0.651 | 5.86 |
| ASD | 0.0304 | 0.991 | 0.0307 | 0.00197 | 0.976 | 8.78 |

**Results for Feature: Filler Word Usage**
Model Fit: $R^2 = 0.067$, Adjusted $R^2 = 0.065$ | Overall Model: $F(6, 2303) = 27.716$, $p = 4.2 \times 10^{-32}$

| Predictor | Coefficient ($\beta$) | Std. Error | t-stat | Effect Size | p-value | p-value (corrected) |
|---|---|---|---|---|---|---|
| (Intercept) | 0.786 | 0.62 | 1.27 | NaN | 0.205 | 1.85 |
| Age | 0.303 | 0.0287 | 10.5 | 0.22 | 2.06E-25 | 1.85E-24 |
| IQ | 0.027 | 0.00499 | 5.41 | 0.113 | 7.09E-08 | 6.38E-07 |
| Sex (Female) | 0.179 | 0.173 | 1.03 | 0.0451 | 0.301 | 2.71 |
| ADHD (Inatt) | -0.0579 | 0.182 | -0.318 | -0.0133 | 0.751 | 6.76 |
| ADHD (Hyper) | 0.105 | 0.194 | 0.538 | 0.0238 | 0.59 | 5.31 |
| ASD | -0.222 | 0.254 | -0.872 | -0.0559 | 0.383 | 3.45 |

**Results for Feature: Self-Focused Speech**
Model Fit: $R^2 = 0.035$, Adjusted $R^2 = 0.032$ | Overall Model: $F(6, 2283) = 13.689$, $p = 2.49 \times 10^{-15}$

| Predictor | Coefficient ($\beta$) | Std. Error | t-stat | Effect Size | p-value | p-value (corrected) |
|---|---|---|---|---|---|---|
| (Intercept) | 10.4 | 2.09 | 4.99 | NaN | 6.54E-07 | 5.89E-06 |
| Age | -0.0261 | 0.0969 | -0.269 | -0.00563 | 0.788 | 7.09 |
| IQ | 0.0797 | 0.0168 | 4.74 | 0.0992 | 2.28E-06 | 2.05E-05 |
| Sex (Female) | -1.04 | 0.585 | -1.78 | -0.0779 | 0.0754 | 0.679 |
| ADHD (Inatt) | 0.415 | 0.615 | 0.674 | 0.0285 | 0.5 | 4.5 |
| ADHD (Hyper) | 0.169 | 0.657 | 0.257 | 0.0114 | 0.797 | 7.17 |
| ASD | 4.5 | 0.858 | 5.24 | 0.337 | 1.74E-07 | 1.57E-06 |

**Results for Feature: % of unique words**
Model Fit: $R^2 = 0.071$, Adjusted $R^2 = 0.068$ | Overall Model: $F(6, 2288) = 29.046$, $p = 1.13 \times 10^{-33}$

| Predictor | Coefficient ($\beta$) | Std. Error | t-stat | Effect Size | p-value | p-value (corrected) |
|---|---|---|---|---|---|---|
| (Intercept) | 88.2 | 0.536 | 165 | NaN | 0 | 0 |
| Age | 0.207 | 0.0248 | 8.35 | 0.175 | 1.19E-16 | 1.07E-15 |
| IQ | 0.0303 | 0.00431 | 7.03 | 0.147 | 2.79E-12 | 2.51E-11 |
| Sex (Female) | 0.389 | 0.15 | 2.6 | 0.114 | 0.00942 | 0.0848 |
| ADHD (Inatt) | -0.26 | 0.157 | -1.65 | -0.0695 | 0.0994 | 0.894 |
| ADHD (Hyper) | -0.255 | 0.168 | -1.52 | -0.0673 | 0.129 | 1.16 |
| ASD | -0.169 | 0.22 | -0.768 | -0.0494 | 0.442 | 3.98 |

**Results for Feature: Lexical Diversity**
Model Fit: $R^2 = 0.238$, Adjusted $R^2 = 0.236$ | Overall Model: $F(6, 2288) = 119.055$, $p = 3.78 \times 10^{-131}$

| Predictor | Coefficient ($\beta$) | Std. Error | t-stat | Effect Size | p-value | p-value (corrected) |
|---|---|---|---|---|---|---|
| (Intercept) | 0.809 | 0.00419 | 193 | NaN | 0 | 0 |
| Age | 0.00439 | 0.000194 | 22.7 | 0.474 | 1.12E-102 | 1.01E-101 |
| IQ | 0.000441 | 3.36E-05 | 13.1 | 0.275 | 4.86E-38 | 4.38E-37 |
| Sex (Female) | 0.00218 | 0.00117 | 1.87 | 0.0817 | 0.062 | 0.558 |
| ADHD (Inatt) | 0.00118 | 0.00123 | 0.962 | 0.0406 | 0.336 | 3.02 |
| ADHD (Hyper) | -0.0023 | 0.00131 | -1.76 | -0.0779 | 0.0792 | 0.713 |
| ASD | -7.41E-05 | 0.00172 | -0.0432 | -0.00277 | 0.966 | 8.69 |

**Results for Feature: Between Word Coherence**
Model Fit: $R^2 = 0.019$, Adjusted $R^2 = 0.017$ | Overall Model: $F(6, 2288) = 7.450$, $p = 6.43 \times 10^{-08}$

| Predictor | Coefficient ($\beta$) | Std. Error | t-stat | Effect Size | p-value | p-value (corrected) |
|---|---|---|---|---|---|---|
| (Intercept) | 0.842 | 0.000925 | 910 | NaN | 0 | 0 |
| Age | 0.00019 | 4.28E-05 | 4.44 | 0.0928 | 9.48E-06 | 8.53E-05 |
| IQ | -1.31E-07 | 7.43E-06 | -0.0177 | -0.00037 | 0.986 | 8.87 |
| Sex (Female) | 0.00026 | 0.000258 | 1.01 | 0.044 | 0.315 | 2.83 |
| ADHD (Inatt) | -0.000329 | 0.000272 | -1.21 | -0.051 | 0.226 | 2.04 |
| ADHD (Hyper) | -0.000165 | 0.00029 | -0.568 | -0.0252 | 0.57 | 5.13 |
| ASD | -0.00143 | 0.000379 | -3.76 | -0.242 | 0.000174 | 0.00157 |

**Results for Feature: Question-Answer Coherence**
Model Fit: $R^2 = 0.011$, Adjusted $R^2 = 0.009$ | Overall Model: $F(6, 2272) = 4.258$, $p = 0.000285$

| Predictor | Coefficient ($\beta$) | Std. Error | t-stat | Effect Size | p-value | p-value (corrected) |
|---|---|---|---|---|---|---|
| (Intercept) | 0.317 | 0.0112 | 28.3 | NaN | 1.37E-151 | 1.23E-150 |
| Age | 0.00134 | 0.000518 | 2.59 | 0.0544 | 0.00958 | 0.0862 |
| IQ | -6.55E-05 | 8.97E-05 | -0.731 | -0.0153 | 0.465 | 4.19 |
| Sex (Female) | 0.0106 | 0.00312 | 3.4 | 0.149 | 0.000695 | 0.00625 |
| ADHD (Inatt) | -0.00173 | 0.00328 | -0.529 | -0.0224 | 0.597 | 5.37 |
| ADHD (Hyper) | 7.30E-05 | 0.0035 | 0.0209 | 0.000928 | 0.983 | 8.85 |
| ASD | -0.00648 | 0.00458 | -1.41 | -0.0913 | 0.158 | 1.42 |

#### Supplementary Section S1.3. Semantic Measures

Detailed model reports for panel e in Figure 3.

**Results for Feature: Factual Memory**
Model Fit: $R^2 = 0.027$, Adjusted $R^2 = 0.024$ | Overall Model: $F(6, 2181) = 10.108$, $p = 4.78 \times 10^{-11}$

| Predictor | Coefficient ($\beta$) | Std. Error | t-stat | Effect Size | p-value | p-value (corrected) |
|---|---|---|---|---|---|---|
| (Intercept) | 1.08 | 0.0604 | 17.8 | NaN | 1.30E-66 | 9.09E-66 |
| Age | 0.00421 | 0.00279 | 1.51 | 0.0323 | 0.131 | 0.92 |
| IQ | 0.00348 | 0.000485 | 7.19 | 0.154 | 8.89E-13 | 6.22E-12 |
| Sex (Female) | -0.00238 | 0.0169 | -0.141 | -0.00634 | 0.888 | 6.21 |
| ADHD (Inatt) | 0.0185 | 0.0177 | 1.04 | 0.0451 | 0.297 | 2.08 |
| ADHD (Hyper) | -0.0275 | 0.0189 | -1.45 | -0.066 | 0.147 | 1.03 |
| ASD | -0.0263 | 0.0245 | -1.07 | -0.0698 | 0.283 | 1.98 |

**Results for Feature: External Emotion Description**
Model Fit: $R^2 = 0.151$, Adjusted $R^2 = 0.149$ | Overall Model: $F(6, 2280) = 67.755$, $p = 8.82 \times 10^{-78}$

| Predictor | Coefficient ($\beta$) | Std. Error | t-stat | Effect Size | p-value | p-value (corrected) |
|---|---|---|---|---|---|---|
| (Intercept) | 1.37 | 0.0486 | 28.1 | NaN | 2.46E-149 | 1.72E-148 |
| Age | -0.0197 | 0.00225 | -8.75 | -0.183 | 4.04E-18 | 2.83E-17 |
| IQ | -0.00096 | 0.00039 | -2.46 | -0.0515 | 0.0139 | 0.0975 |
| Sex (Female) | -0.0073 | 0.0136 | -0.539 | -0.0236 | 0.59 | 4.13 |
| ADHD (Inatt) | 0.0178 | 0.0143 | 1.25 | 0.0526 | 0.212 | 1.49 |
| ADHD (Hyper) | 0.00984 | 0.0152 | 0.647 | 0.0287 | 0.518 | 3.63 |
| ASD | -0.00095 | 0.02 | -0.0476 | -0.00307 | 0.962 | 6.73 |

**Results for Feature: Self Emotion Description**
Model Fit: $R^2 = 0.039$, Adjusted $R^2 = 0.036$ | Overall Model: $F(6, 2259) = 15.254$, $p = 3.34 \times 10^{-17}$

| Predictor | Coefficient ($\beta$) | Std. Error | t-stat | Effect Size | p-value | p-value (corrected) |
|---|---|---|---|---|---|---|
| (Intercept) | 1.16 | 0.0378 | 30.7 | NaN | 4.92E-173 | 3.44E-172 |
| Age | -0.0051 | 0.00175 | -2.92 | -0.0614 | 0.00355 | 0.0248 |
| IQ | -0.00154 | 0.000304 | -5.05 | -0.106 | 4.67E-07 | 3.27E-06 |
| Sex (Female) | 0.0421 | 0.0106 | 3.98 | 0.176 | 7.00E-05 | 0.000490 |
| ADHD (Inatt) | 0.00258 | 0.0111 | 0.232 | 0.00984 | 0.816 | 5.72 |
| ADHD (Hyper) | 0.0166 | 0.0118 | 1.4 | 0.0625 | 0.161 | 1.13 |
| ASD | -0.0663 | 0.0156 | -4.25 | -0.277 | 2.22E-05 | 0.000155 |

**Results for Feature: Narrative Detail**
Model Fit: $R^2 = 0.190$, Adjusted $R^2 = 0.187$ | Overall Model: $F(6, 2254) = 87.911$, $p = 2.79 \times 10^{-99}$

| Predictor | Coefficient ($\beta$) | Std. Error | t-stat | Effect Size | p-value | p-value (corrected) |
|---|---|---|---|---|---|---|
| (Intercept) | 0.747 | 0.04 | 18.7 | NaN | 1.14E-72 | 7.99E-72 |
| Age | 0.0353 | 0.00185 | 19.1 | 0.402 | 2.96E-75 | 2.08E-74 |
| IQ | 0.00295 | 0.00032 | 9.22 | 0.194 | 6.49E-20 | 4.54E-19 |
| Sex (Female) | 0.0427 | 0.0111 | 3.84 | 0.169 | 0.000128 | 0.000896 |
| ADHD (Inatt) | 0.0196 | 0.0117 | 1.67 | 0.0711 | 0.0941 | 0.659 |
| ADHD (Hyper) | -0.0306 | 0.0125 | -2.45 | -0.11 | 0.0143 | 0.1 |
| ASD | -0.0904 | 0.0164 | -5.52 | -0.358 | 3.76E-08 | 2.63E-07 |

**Results for Feature: Thematic Understanding**
Model Fit: $R^2 = 0.160$, Adjusted $R^2 = 0.158$ | Overall Model: $F(6, 2017) = 64.100$, $p = 4.81 \times 10^{-73}$

| Predictor | Coefficient ($\beta$) | Std. Error | t-stat | Effect Size | p-value | p-value (corrected) |
|---|---|---|---|---|---|---|
| (Intercept) | 0.0481 | 0.0854 | 0.563 | NaN | 0.573 | 4.01 |
| Age | 0.0652 | 0.00389 | 16.8 | 0.373 | 3.85E-59 | 2.70E-58 |
| IQ | 0.00605 | 0.000681 | 8.89 | 0.198 | 1.34E-18 | 9.39E-18 |
| Sex (Female) | -0.0101 | 0.0233 | -0.435 | -0.0203 | 0.663 | 4.64 |
| ADHD (Inatt) | 0.016 | 0.0247 | 0.649 | 0.0292 | 0.516 | 3.62 |
| ADHD (Hyper) | -0.0357 | 0.0264 | -1.35 | -0.0642 | 0.176 | 1.23 |
| ASD | -0.165 | 0.0346 | -4.78 | -0.33 | 1.90E-06 | 1.33E-05 |

**Results for Feature: Positive Preferences**
Model Fit: $R^2 = 0.047$, Adjusted $R^2 = 0.044$ | Overall Model: $F(6, 2015) = 16.458$, $p = 1.35 \times 10^{-18}$

| Predictor | Coefficient ($\beta$) | Std. Error | t-stat | Effect Size | p-value | p-value (corrected) |
|---|---|---|---|---|---|---|
| (Intercept) | 0.833 | 0.0375 | 22.3 | NaN | 2.90E-98 | 2.03E-97 |
| Age | 0.00486 | 0.00171 | 2.84 | 0.0633 | 0.00452 | 0.0316 |
| IQ | 0.000754 | 0.0003 | 2.51 | 0.0559 | 0.0121 | 0.085 |
| Sex (Female) | 0.0544 | 0.0105 | 5.19 | 0.242 | 2.31E-07 | 1.62E-06 |
| ADHD (Inatt) | -0.0112 | 0.0111 | -1.01 | -0.0454 | 0.313 | 2.19 |
| ADHD (Hyper) | -0.0145 | 0.0118 | -1.23 | -0.0579 | 0.22 | 1.54 |
| ASD | -0.0792 | 0.0154 | -5.15 | -0.352 | 2.82E-07 | 1.97E-06 |

**Results for Feature: Negative Preferences**
Model Fit: $R^2 = 0.019$, Adjusted $R^2 = 0.016$ | Overall Model: $F(6, 1939) = 6.323$, $p = 1.34 \times 10^{-06}$

| Predictor | Coefficient ($\beta$) | Std. Error | t-stat | Effect Size | p-value | p-value (corrected) |
|---|---|---|---|---|---|---|
| (Intercept) | 0.826 | 0.0341 | 24.2 | NaN | 3.01E-113 | 2.10E-112 |
| Age | 0.00438 | 0.00155 | 2.82 | 0.0642 | 0.00478 | 0.0334 |
| IQ | 6.19E-05 | 0.000273 | 0.227 | 0.00515 | 0.821 | 5.74 |
| Sex (Female) | 0.0361 | 0.00947 | 3.81 | 0.181 | 0.000145 | 0.00102 |
| ADHD (Inatt) | 0.000497 | 0.01 | 0.0496 | 0.00228 | 0.96 | 6.72 |
| ADHD (Hyper) | -0.00878 | 0.0107 | -0.822 | -0.0396 | 0.411 | 2.88 |
| ASD | -0.0353 | 0.0138 | -2.55 | -0.177 | 0.0108 | 0.0755 |

#### Supplementary Section S1.4. Vocal Prosody Measures

Detailed model reports for panel b in Figure 4.

**Results for Feature: Pitch**
Model Fit: $R^2 = 0.160$, Adjusted $R^2 = 0.158$ | Overall Model: $F(6, 2291) = 72.592$, $p = 4.38 \times 10^{-83}$

| Predictor | Coefficient ($\beta$) | Std. Error | t-stat | Effect Size | p-value | p-value (corrected) |
|---|---|---|---|---|---|---|
| (Intercept) | 238 | 6.63 | 36 | NaN | 7.62E-225 | 4.57E-224 |
| Age | -5.84 | 0.307 | -19 | -0.398 | 3.80E-75 | 2.28E-74 |
| IQ | 0.0328 | 0.0533 | 0.614 | 0.0128 | 0.539 | 3.23 |
| Sex (Female) | 5.12 | 1.85 | 2.77 | 0.121 | 0.00569 | 0.0341 |
| ADHD (Inatt) | 1.79 | 1.94 | 0.922 | 0.0389 | 0.356 | 2.14 |
| ADHD (Hyper) | 5.32 | 2.08 | 2.56 | 0.113 | 0.0105 | 0.063 |
| ASD | 11.1 | 2.72 | 4.07 | 0.262 | 4.79E-05 | 0.000287 |

**Results for Feature: Pitch Variation**
Model Fit: $R^2 = 0.171$, Adjusted $R^2 = 0.169$ | Overall Model: $F(6, 2291) = 78.700$, $p = 1.14 \times 10^{-89}$

| Predictor | Coefficient ($\beta$) | Std. Error | t-stat | Effect Size | p-value | p-value (corrected) |
|---|---|---|---|---|---|---|
| (Intercept) | 77.8 | 2.64 | 29.4 | NaN | 1.13E-161 | 6.79E-161 |
| Age | -2.34 | 0.122 | -19.1 | -0.399 | 9.72E-76 | 5.83E-75 |
| IQ | 0.00506 | 0.0213 | 0.238 | 0.00497 | 0.812 | 4.87 |
| Sex (Female) | 1.9 | 0.738 | 2.57 | 0.112 | 0.0102 | 0.0615 |
| ADHD (Inatt) | 0.312 | 0.775 | 0.403 | 0.017 | 0.687 | 4.12 |
| ADHD (Hyper) | 3.03 | 0.828 | 3.66 | 0.162 | 0.000263 | 0.00158 |
| ASD | 5.67 | 1.08 | 5.23 | 0.336 | 1.81E-07 | 1.09E-06 |

**Results for Feature: Loudness**
Model Fit: $R^2 = 0.030$, Adjusted $R^2 = 0.028$ | Overall Model: $F(6, 2291) = 11.905$, $p = 3.38 \times 10^{-13}$

| Predictor | Coefficient ($\beta$) | Std. Error | t-stat | Effect Size | p-value | p-value (corrected) |
|---|---|---|---|---|---|---|
| (Intercept) | 57.3 | 0.956 | 59.9 | NaN | 0 | 0 |
| Age | 0.0684 | 0.0442 | 1.55 | 0.0323 | 0.122 | 0.733 |
| IQ | -0.0149 | 0.00769 | -1.94 | -0.0405 | 0.0526 | 0.316 |
| Sex (Female) | -0.839 | 0.267 | -3.14 | -0.137 | 0.00169 | 0.0101 |
| ADHD (Inatt) | -0.559 | 0.28 | -1.99 | -0.084 | 0.0462 | 0.277 |
| ADHD (Hyper) | 1.29 | 0.299 | 4.32 | 0.191 | 1.63E-05 | 9.80E-05 |
| ASD | 1.87 | 0.392 | 4.78 | 0.307 | 1.88E-06 | 1.13E-05 |

**Results for Feature: Loudness Variation**
Model Fit: $R^2 = 0.026$, Adjusted $R^2 = 0.024$ | Overall Model: $F(6, 2291) = 10.372$, $p = 2.28 \times 10^{-11}$

| Predictor | Coefficient ($\beta$) | Std. Error | t-stat | Effect Size | p-value | p-value (corrected) |
|---|---|---|---|---|---|---|
| (Intercept) | 6.45 | 0.3 | 21.5 | NaN | 9.84E-94 | 5.91E-93 |
| Age | -0.0788 | 0.0139 | -5.68 | -0.119 | 1.48E-08 | 8.89E-08 |
| IQ | -0.0013 | 0.00241 | -0.54 | -0.0113 | 0.589 | 3.54 |
| Sex (Female) | -0.3 | 0.0836 | -3.59 | -0.157 | 0.000335 | 0.00201 |
| ADHD (Inatt) | 0.0441 | 0.0878 | 0.502 | 0.0211 | 0.616 | 3.69 |
| ADHD (Hyper) | 0.0336 | 0.0938 | 0.358 | 0.0159 | 0.72 | 4.32 |
| ASD | 0.323 | 0.123 | 2.63 | 0.169 | 0.00865 | 0.0519 |

**Results for Feature: Dysphonia**
Model Fit: $R^2 = 0.066$, Adjusted $R^2 = 0.064$ | Overall Model: $F(6, 2291) = 27.179$, $p = 1.84 \times 10^{-31}$

| Predictor | Coefficient ($\beta$) | Std. Error | t-stat | Effect Size | p-value | p-value (corrected) |
|---|---|---|---|---|---|---|
| (Intercept) | 15.4 | 0.0683 | 225 | NaN | 0 | 0 |
| Age | 0.0148 | 0.00316 | 4.7 | 0.0982 | 2.75E-06 | 1.65E-05 |
| IQ | -0.000102 | 0.000549 | -0.186 | -0.00388 | 0.853 | 5.12 |
| Sex (Female) | -0.0269 | 0.019 | -1.41 | -0.0619 | 0.157 | 0.944 |
| ADHD (Inatt) | -0.00671 | 0.02 | -0.336 | -0.0141 | 0.737 | 4.42 |
| ADHD (Hyper) | 0.0301 | 0.0214 | 1.41 | 0.0623 | 0.16 | 0.958 |
| ASD | 0.222 | 0.028 | 7.92 | 0.509 | 3.54E-15 | 2.12E-14 |

**Results for Feature: Breathiness**
Model Fit: $R^2 = 0.034$, Adjusted $R^2 = 0.032$ | Overall Model: $F(6, 2291) = 13.452$, $p = 4.77 \times 10^{-15}$

| Predictor | Coefficient ($\beta$) | Std. Error | t-stat | Effect Size | p-value | p-value (corrected) |
|---|---|---|---|---|---|---|
| (Intercept) | 0.732 | 0.0106 | 69.4 | NaN | 0 | 0 |
| Age | -0.00234 | 0.000488 | -4.78 | -0.1 | 1.83E-06 | 1.10E-05 |
| IQ | -8.83E-05 | 8.48E-05 | -1.04 | -0.0218 | 0.298 | 1.79 |
| Sex (Female) | 0.00525 | 0.00294 | 1.78 | 0.078 | 0.0747 | 0.448 |
| ADHD (Inatt) | -0.00426 | 0.00309 | -1.38 | -0.058 | 0.169 | 1.01 |
| ADHD (Hyper) | 0.00585 | 0.0033 | 1.77 | 0.0785 | 0.0764 | 0.458 |
| ASD | 0.028 | 0.00432 | 6.47 | 0.416 | 1.17E-10 | 7.02E-10 |

#### Supplementary Section S1.5. Movement Measures

Detailed model reports for panel b in Figure 5.

**Results for Feature: Face**
Model Fit: $R^2 = 0.107$, Adjusted $R^2 = 0.105$ | Overall Model: $F(6, 2320) = 46.543$, $p = 4.35 \times 10^{-54}$

| Predictor | Coefficient ($\beta$) | Std. Error | t-stat | Effect Size | p-value | p-value (corrected) |
|---|---|---|---|---|---|---|
| (Intercept) | -3.4 | 0.0721 | -47.1 | NaN | 0 | 0 |
| Age | -0.0443 | 0.00334 | -13.2 | -0.275 | 1.23E-38 | 4.91E-38 |
| IQ | 0.000891 | 0.000579 | 1.54 | 0.0319 | 0.124 | 0.497 |
| Sex (Female) | -0.0102 | 0.0201 | -0.506 | -0.022 | 0.613 | 2.45 |
| ADHD (Inatt) | -0.0182 | 0.0212 | -0.858 | -0.0359 | 0.391 | 1.56 |
| ADHD (Hyper) | 0.134 | 0.0227 | 5.89 | 0.26 | 4.35E-09 | 1.74E-08 |
| ASD | 0.0592 | 0.0295 | 2.01 | 0.128 | 0.0446 | 0.178 |

**Results for Feature: Eye**
Model Fit: $R^2 = 0.110$, Adjusted $R^2 = 0.107$ | Overall Model: $F(6, 2320) = 47.590$, $p = 2.75 \times 10^{-55}$

| Predictor | Coefficient ($\beta$) | Std. Error | t-stat | Effect Size | p-value | p-value (corrected) |
|---|---|---|---|---|---|---|
| (Intercept) | -3.73 | 0.0741 | -50.2 | NaN | 0 | 0 |
| Age | -0.0458 | 0.00344 | -13.3 | -0.277 | 3.46E-39 | 1.38E-38 |
| IQ | 0.0011 | 0.000595 | 1.85 | 0.0384 | 0.0646 | 0.259 |
| Sex (Female) | -0.0346 | 0.0207 | -1.67 | -0.0726 | 0.0942 | 0.377 |
| ADHD (Inatt) | -0.0223 | 0.0218 | -1.03 | -0.0429 | 0.305 | 1.22 |
| ADHD (Hyper) | 0.13 | 0.0233 | 5.57 | 0.246 | 2.82E-08 | 1.13E-07 |
| ASD | 0.0612 | 0.0303 | 2.02 | 0.128 | 0.0434 | 0.174 |

**Results for Feature: Mouth**
Model Fit: $R^2 = 0.104$, Adjusted $R^2 = 0.102$ | Overall Model: $F(6, 2320) = 44.884$, $p = 3.49 \times 10^{-52}$

| Predictor | Coefficient ($\beta$) | Std. Error | t-stat | Effect Size | p-value | p-value (corrected) |
|---|---|---|---|---|---|---|
| (Intercept) | -2.77 | 0.0692 | -40 | NaN | 5.29E-267 | 2.12E-266 |
| Age | -0.0416 | 0.00321 | -13 | -0.27 | 2.91E-37 | 1.16E-36 |
| IQ | 0.000733 | 0.000556 | 1.32 | 0.0274 | 0.187 | 0.749 |
| Sex (Female) | 0.00479 | 0.0193 | 0.248 | 0.0108 | 0.804 | 3.22 |
| ADHD (Inatt) | -0.0146 | 0.0203 | -0.718 | -0.03 | 0.473 | 1.89 |
| ADHD (Hyper) | 0.131 | 0.0218 | 6.04 | 0.266 | 1.76E-09 | 7.03E-09 |
| ASD | 0.058 | 0.0283 | 2.05 | 0.13 | 0.0406 | 0.162 |

**Results for Feature: Upper Body**
Model Fit: $R^2 = 0.180$, Adjusted $R^2 = 0.178$ | Overall Model: $F(6, 2334) = 85.393$, $p = 5.83 \times 10^{-97}$

| Predictor | Coefficient ($\beta$) | Std. Error | t-stat | Effect Size | p-value | p-value (corrected) |
|---|---|---|---|---|---|---|
| (Intercept) | -1.25 | 0.0639 | -19.6 | NaN | 3.69E-79 | 1.48E-78 |
| Age | -0.0601 | 0.00296 | -20.3 | -0.419 | 3.19E-84 | 1.28E-83 |
| IQ | -0.0024 | 0.000513 | -4.68 | -0.0968 | 3.06E-06 | 1.22E-05 |
| Sex (Female) | 0.0861 | 0.0178 | 4.83 | 0.209 | 1.47E-06 | 5.90E-06 |
| ADHD (Inatt) | 0.00457 | 0.0188 | 0.244 | 0.0102 | 0.808 | 3.23 |
| ADHD (Hyper) | 0.0888 | 0.0201 | 4.42 | 0.194 | 1.02E-05 | 4.08E-05 |
| ASD | 0.0441 | 0.0262 | 1.68 | 0.107 | 0.0925 | 0.37 |

#### Supplementary Section S2. Diarizing Prompt for LLM

```
You are given a transcript of an interview between a clinician (interviewer) and a
subject. The interviewer asks questions from a predefined list, though slight
variations may occur. Your task is to label the dialogue as either "Interviewer:"
or "Subject:", removing timestamps while preserving the dialogue structure.

### Instructions:
1. Identify lines spoken by the **interviewer** based on a predefined question list.
   These questions may have slight variations but follow the same general intent.
2. Label all other responses as **"Subject:"**
3. Remove timestamps but keep the text structure intact.
4. Ensure proper formatting with a new line between each labeled turn.

### Example Input:
0.709 -> 2.83: I hope you enjoyed the last movie about the puppy.
2.95 -> 3.851: Have you seen it before?
4.451 -> 4.751: No.
5.231 -> 6.792: Can you tell me what happened in the movie?
6.852 -> 8.013: Try to tell the whole story.
8.093 -> 11.515: Remember that stories have a beginning, things that happen, and an ending.
12.895 -> 15.877: I didn't watch the whole thing though.
16.637 -> 18.478: Can you tell me about the part that you did watch?
19.139 -> 19.639: Okay.

### Example Output:
Interviewer: I hope you enjoyed the last movie about the puppy.
Interviewer: Have you seen it before?
Subject: No.
Interviewer: Can you tell me what happened in the movie?
Interviewer: Try to tell the whole story.
Interviewer: Remember that stories have a beginning, things that happen, and an ending.
Subject: I didn't watch the whole thing though.
Interviewer: Can you tell me about the part that you did watch?
Subject: Okay.

### If the transcript contains these specific sentences, you will label them as "clip"
instead of interviewer or subject:
Whoa! Cool!
You gotta be kidding me
Get lost
Mom, I'll/we'll be outside

### The interviewer question list:
So I hope you enjoyed the last movie. Have you seen it before?
So can you tell me what happened in the movie? Try to tell the whole story.
Remember that stories have a beginning, things that happen, and an ending.
Do you remember anything else from the story?
So what are some of the things you liked about the movie?
What are some of the things you didn't like about the movie?
Who gave the boy a box?
What was in the box?
What was the boy doing before he got the box?
What was the puppy playing with?
How are the puppy and the boy the same?
So in the movie, who is missing a leg? The boy, the puppy, both the boy and the puppy,
or no one?
```

```
So we're going to watch a short clip from the movie and then we'll talk about it.
How do you think the puppy was feeling?
How do you think the boy was feeling?
And how did you feel while you were watching that part?
How do you think the puppy was feeling?
How do you think the boy was feeling?
And how did you feel while you were watching that part?
How do you think the puppy was feeling?
How do you think the boy was feeling?
And how did you feel while you were watching that part?
How do you think the puppy was feeling?
How do you think the boy was feeling?
And how did you feel while you were watching this part?
Great, thank you

### In some cases, the interviewer may ask additional questions or make comments.
You should label these as "Interviewer:" as well using their specific context.

### The transcript:
<<transcript>>
```

### Supplementary Section S3. Question-Answer Extraction Prompt for LLM

```
Task: Extract Answers from Interview Transcript

You are tasked with extracting the participant’s answers to a specific set of predefined
questions from an interview transcript. The questions relate to the short film
"The Present." The interviewer *may* have asked these questions verbatim, asked them
in a slightly different way, or not asked them at all. The subject may have answered
directly, indirectly, partially, over multiple turns, or not at all.
In the transcript the interviewer asks these eleven questions in order:
1.  "I hope you enjoyed the last movie. Have you seen it before?"
2.  "Can you tell me what happened in the movie? Try to tell the whole story. Remember
    that stories have a beginning, things that happen, and an ending."
3.  "Do you remember anything else from the story?"
4.  "What are some of the things you liked about the movie?"
5.  "What are some of the things you didn’t like about the movie?"
6.  "Who gave the boy a box?"
7.  "What was in the box?"
8.  "What was the boy doing before he got the box?"
9.  "What was the puppy playing with?"
10. "How are the puppy and the boy the same?"
11. "In the movie, who is missing a leg? The boy, the puppy, both the boy and the puppy,
    or no one?"

After these questions, the interviewer then shows the subject four short clips from
the movie and asks the subject to describe how the characters are feeling in each clip.
The interviewer may say something like:
- "Let’s watch a short clip from the movie, and then we’ll talk about it."
- "Now let’s watch another clip"
- "One more clip"
- "One last clip"

After watching each clip, the interviewer asks the subject how they think the characters
are feeling and how they themselves feel while watching the clip. The interviewer may ask
these questions in various ways, such as:
- "How do you think the puppy was feeling?"
- "How do you think the boy was feeling?"
- "And how did you feel while you were watching that part?"

Input:
* A transcript of a conversation between an interviewer and a subject discussing the short
  film "The Present."

Output:
Your output will consist of two parts:

**Part 1: Question-Answer Extraction**

A series of lines, one for each predefined question. Each line will follow this format:
‘"Question" -> "Answer"‘

Where:
* ‘"Question"‘ is one of the predefined questions listed below (use the *exact* wording
  provided here, even if the interviewer phrased it differently).
* ‘"Answer"‘ is the subject’s answer to the question.
  * If the question was answered (directly or indirectly), provide the subject’s complete
```

```
    answer, concatenating all relevant parts of their response, even if it spans multiple
    turns or requires piecing together information from different parts of the transcript.
  * If the question was *not* asked, or if the subject provided *no* discernible answer
    (even after considering the surrounding context), leave the "Answer" field *blank*.
    Do *not* write "N/A," "No answer," or any other placeholder. Just leave it blank.
  * If parts of different answers are mixed, extract only the answer to the intended
    question.

Predefined Questions:
1.  "I hope you enjoyed the last movie. Have you seen it before?"
2.  "Can you tell me what happened in the movie? Try to tell the whole story. Remember
    that stories have a beginning, things that happen, and an ending."
3.  "Do you remember anything else from the story?"
4.  "What are some of the things you liked about the movie?"
5.  "What are some of the things you didn't like about the movie?"
6.  "Who gave the boy a box?"
7.  "What was in the box?"
8.  "What was the boy doing before he got the box?"
9.  "What was the puppy playing with?"
10. "How are the puppy and the boy the same?"
11. "In the movie, who is missing a leg? The boy, the puppy, both the boy and the puppy,
    or no one?"
12. "How do you think the puppy was feeling? (after watching first clip)"
13. "How do you think the boy was feeling? (after watching first clip)"
14. "And how did you feel while you were watching that part? (after watching first clip)"
15. "How do you think the puppy was feeling? (after watching second clip)"
16. "How do you think the boy was feeling? (after watching second clip)"
17. "And how did you feel while you were watching that part? (after watching second clip)"
18. "How do you think the puppy was feeling? (after watching third clip)"
19. "How do you think the boy was feeling? (after watching third clip)"
20. "And how did you feel while you were watching that part? (after watching third clip)"
21. "How do you think the puppy was feeling? (after watching fourth clip)"
22. "How do you think the boy was feeling? (after watching fourth clip)"
23. "And how did you feel while you were watching that part? (after watching fourth clip)"

**Part 2: Conversation Completion Rating**

A single line in the following format:
'"Conversation Completion Rating" -> "Rating"'

Where:
* '"Rating"' is a numerical score between 0 and 1 (inclusive) representing the overall
  completion of the conversation, *specifically with respect to the predefined questions*.
  * **1:** Represents a perfect completion, where all of the predefined questions were
    asked (or closely paraphrased), and the subject provided clear and complete answers
    to each.
  * **0:** Represents a very poor completion, where many questions were not asked, or the
    subject consistently failed to answer the questions that were asked.
  * **Intermediate values:** Represent varying degrees of completion quality. A score
    of 0.5, for example, would suggest that roughly half of the questions were asked and
    answered reasonably well. Use your judgment to assign a score that reflects the
    overall completeness and clarity of the question-answer exchange related to the
    *predefined questions*. Do *not* assess the overall quality of the conversation
    beyond its adherence to addressing the core content of these questions.

Instructions:

1.  **Read the Entire Transcript:** Carefully read the entire transcript to understand the
```

```
    completion of the conversation and the context of each statement.
2.  **Identify Questions and Answers:** For each predefined question:
    * **Search for the Question:** Look for the exact question, or a close paraphrase, in
      the interviewer's speech. Note that the interviewer might rephrase, prompt, or
      interrupt the subject.
    * **Locate the Answer (If Present):** If the question (or a close variant) was asked,
      carefully examine the subject's subsequent responses. The answer might be:
      * **Direct and Immediate:** Right after the question.
      * **Indirect:** Implied by their statements, requiring inference.
      * **Partial:** Only part of the answer is given directly.
      * **Scattered:** Pieces of the answer are given across multiple turns, possibly
        with interviewer prompts or interruptions.
      * **Non-existent:** The subject might not answer at all.
    * **Use Context:** Consider surrounding conversational turns from both interviewer and
      subject to determine if a particular utterance constitutes an answer, even an indirect
      or partial one.
    * **Combine Response Parts:** If the answer is spread over multiple turns, concatenate
      *all relevant parts* of the subject's response into a single, coherent answer.
      Remove any interviewer interjections or prompts from within the concatenated answer.
      Only include the *subject's* words in the "Answer" field.
    * **For the questions after the clips:** Because there are multiple clips, you will
      need to extract the answers to these questions from the conversation after each clip
      is shown. The interviewer will ask the subject how they think the characters are
      feeling and how they themselves feel while watching the clip. However, the order of
      the clips is always the same. Sometimes part of the dialogue is present on the
      transcript, so in those cases, use that as clue to know to what clip the question
      is referring to. If you can't clearly know what question/answer corresponds to what
      clip, restrain from returning an answer for that specific question, leave the
      "Answer" field blank.:
      * **Clip 1:** the kid opens the present and sees the puppy, the transcript might
        say "Whoa, cool."
      * **Clip 2:** the kid throws the puppy away, the transcript might say "You've got
        to be kidding me."
      * **Clip 3:** the kid kicks the puppy, the transcript might say "Get lost!"
      * **Clip 4:** the kid plays with the puppy, the transcript might say "Mom, I'll /
        We'll be outside."
3.  **Output (Part 1):** Generate the question-answer extraction in the specified
    '"Question" -> "Answer"' format. Leave the "Answer" field blank if the question was
    not asked or if no answer can be found.
4.  **Assess Conversation Completion (Part 2):** After extracting the question-answer
    pairs, evaluate the *overall completion* of the conversation *with respect to the
    predefined questions*. Consider:
    * How many of the predefined questions (or close paraphrases) were asked by the
      interviewer?
    * Did the conversation stay focused on the topics covered by the predefined questions,
      or did it frequently deviate?
5.  **Output (Part 2):** Provide the "Conversation Completion Rating" on a separate line,
    using a number between 0 and 1.

Example 1:

    (example 1 transcript):
        Interviewer: So I hope you enjoyed the last movie.
        Interviewer: Have you seen it before?
        Subject: I have at the mock MRI.
        Interviewer: Oh, cool.
        Subject: I liked it, though.
        Interviewer: Great.
```

```
Interviewer: So can you tell me what happened in the movie?
Interviewer: Try to tell the whole story.
Interviewer: Remember that stories have a beginning, things that happen, and an ending.
Subject: So there's a boy who was disabled, and he didn't want to go outside.
Subject: He wasn't in the mood for anything except for his video games.
Subject: and then like his mom was trying to like help him go like get some fresh air
         and then but like everything like his mom tried to do didn't work so then he
         then she bought um him a puppy so then he started like he didn't want like
         to see the puppy like anymore because then he realized he realized that he
         was also like disabled
Subject: But then he started having a liking to him because he felt like the puppy
         and him were like the same.
Subject: So he started playing with the puppy more.
Subject: So then they went outside and they started playing with the ball.
Interviewer: Great.
Interviewer: Do you remember anything else from the story?
Subject: I think that's much it.
Interviewer: Great.
Interviewer: So what are some of the things you liked about the movie?
Subject: I liked how he had a good relationship with his doll.
Interviewer: What are some of the things you didn't like about the movie?
Subject: I liked everything about the movie, but
Subject: I was hoping it would be longer.
Interviewer: Who gave the boy a box?
Subject: His mother.
Interviewer: What was in the box?
Subject: The puppy.
Interviewer: What was the boy doing before he got the box?
Subject: He wasn't really paying attention to anything.
Subject: He was glued to his video games.
Interviewer: What was the puppy playing with?
Subject: He was playing with the red rubber ball.
Interviewer: How are the puppy and the boy the same?
Subject: Both of them each have a disabled part of their body.
Interviewer: So in the movie, who is missing a leg?
Interviewer: The boy, the puppy, both the boy and the puppy, or no one?
Subject: The boy.
Interviewer: Great.
Interviewer: So we're going to watch a short clip from the movie and then we'll talk
             about it.
Subject: Okay.
clip: Whoa, cool.
Interviewer: How do you think the puppy was feeling?
Subject: Happy.
Interviewer: How do you think the boy was feeling?
Subject: He's feeling happy too.
Interviewer: And how did you feel while you were watching that pot?
Subject: I guess happy.
Interviewer: Right.
Interviewer: How do you think the puppy was feeling?
Subject: Sad.
Interviewer: How do you think the boy was feeling?
Subject: He was feeling angry.
Interviewer: And how did you feel while you were watching that mom?
Subject: I didn't like it.
Interviewer: How do you think the puppy was feeling?
Subject: He didn't mind.
Interviewer: How do you think the boy was feeling?
```

```
        Subject: He was still angry.
        Interviewer: And how did you feel while you were watching that pup?
        Subject: I feel bad for the puppy.
        Interviewer: Great.
        clip: Mom, I'll be outside.
        Interviewer: How do you think the puppy was feeling?
        Subject: Um, happy.
        Interviewer: How do you think the boy was feeling?
        Subject: He was feeling happy too.
        Interviewer: And how did you feel while you were watching this pot?
        Subject: I was happy.
        Interviewer: Great, thank you.

    (example 1 output):
        "I hope you enjoyed the last movie. Have you seen it before?" -> "I have at the mug MRI."
        "Can you tell me what happened in the movie? Try to tell the whole story. Remember
        that stories have a beginning, things that happen, and an ending." -> "So there's a boy
        who was disabled, and he didn't want to go outside. He wasn't in the mood for anything
        except for his video games. And then like his mom was trying to like help him go like
        get some fresh air and then but like everything like his mom tried to do didn't work so
        then he then she bought um him a puppy so then he started like he didn't want like to
        see the puppy like anymore because then he realized he realized that he was also like
        disabled. But then he started having a liking to him because he felt like the puppy and
        him were like the same. So he started playing with the puppy more. So then they went
        outside and they started playing with the ball."
        "Do you remember anything else from the story?" -> "I think that's much it."
        "What are some of the things you liked about the movie?" -> "I liked how he had a good
        relationship with his doll."
        "What are some of the things you didn't like about the movie?" -> "I liked everything
        about the movie, but I was hoping it would be longer."
        "Who gave the boy a box" -> "His mother."
        "What was in the box?" -> "The puppy."
        "What was the boy doing before he got the box?" -> "He wasn't really paying attention to
        anything. He was glued to his video games."
        "What was the puppy playing with?" -> "He was playing with the red rubber ball."
        "How are the puppy and the boy the same?" -> "Both of them each have a disabled part of
        their body."
        "In the movie, who is missing a leg? The boy, the puppy, both the boy and the puppy, or
        no one?" ->  "The boy."
        "How do you think the puppy was feeling? (after watching first clip)" -> "Happy."
        "How do you think the boy was feeling? (after watching first clip)" -> "He's feeling
        happy too."
        "And how did you feel while you were watching that part? (after watching first clip)"
        -> "I guess happy."
        "How do you think the puppy was feeling? (after watching second clip)" -> "Sad."
        "How do you think the boy was feeling? (after watching second clip)" -> "He was feeling
        angry."
        "And how did you feel while you were watching that part? (after watching second clip)"
        -> "I didn't like it."
        "How do you think the puppy was feeling? (after watching third clip)" -> "He didn't mind."
        "How do you think the boy was feeling? (after watching third clip)" -> "He was still angry."
        "And how did you feel while you were watching that part? (after watching third clip)"
        -> "I feel bad for the puppy"
        "How do you think the puppy was feeling? (after watching fourth clip)" -> "Um, happy."
        "How do you think the boy was feeling? (after watching fourth clip)" -> "He was feeling
        happy too."
        "And how did you feel while you were watching that part? (after watching fourth clip)"
        -> "I was happy."
```

```
        "Rating" -> "1"

Example 2:

    (example 2 transcript):
        Interviewer: Okay, so I hope you enjoyed the last movie.
        Interviewer: Have you seen it before?
        Subject: No
        Interviewer: Um, so I'm going to need you to talk a little bit.
        Interviewer: Is that okay?
        Subject: Okay.
        Interviewer: Can you tell me what happened in the movie?
        Interviewer: Try to tell the whole story.
        Interviewer: Remember that stories have a beginning, things that happen, and an ending.
        Interviewer: Great.
        Interviewer: Do you remember anything else from the story?
        Interviewer: Okay.
        Interviewer: What are some of the things you liked about the movie?
        Interviewer: And what are some of the things you didn't like about the movie?
        Interviewer: So who gave the boy a box?
        Interviewer: What was in the box?
        Interviewer: What was the boy doing before he got the box?
        Interviewer: What was the puppy playing with?
        Interviewer: How are the puppy and the boy the same?
        Interviewer: Perfect.
        Interviewer: So in the movie, who is missing a leg?
        Interviewer: The boy, the puppy, both the boy and the puppy, or no one?
        Subject: Both the boy and the puppy.
        Interviewer: Great.
        Interviewer: So we're going to watch a couple of short clips from the movie and
                     talk about them, OK?
        Subject: OK.
        clip: Whoa.
        clip: Cool.
        Interviewer: How do you think the puppy was feeling?
        Subject: Happy.
        Interviewer: How do you think the boy was feeling?
        Subject: Happy.
        Interviewer: Happy?
        Interviewer: And how did you feel while you were watching this part?
        Subject: Good.
        Interviewer: Good?
        Interviewer: Great.
        clip: You've got to be kidding me.
        Interviewer: How do you think the puppy was feeling?
        Interviewer: And how do you think the boy was feeling?
        Interviewer: And how did you feel while you were watching this part?
        clip: Get lost!
        Interviewer: How do you think the puppy was feeling?
        Interviewer: And how do you think the boy was feeling?
        Interviewer: And how did you feel while you were watching this pot?
        Interviewer: Okay, one more.
        clip: Mom!
        clip: We'll be outside!
        Interviewer: How do you think the puppy was feeling?
        Subject: Happy.
        Interviewer: How do you think the boy was feeling?
        Subject: Happy.
```

```
        Interviewer: And how did you feel while you were watching his paw?
        Subject: Happy.
        Interviewer: Great, thank you.


    (example 2 output):
        "I hope you enjoyed the last movie. Have you seen it before?" -> "No."
        "Can you tell me what happened in the movie? Try to tell the whole story. Remember
        that stories have a beginning, things that happen, and an ending." -> ""
        "Do you remember anything else from the story?" -> ""
        "What are some of the things you liked about the movie?" -> ""
        "What are some of the things you didn't like about the movie?" -> ""
        "Who gave the boy a box" -> ""
        "What was in the box?" -> ""
        "What was the boy doing before he got the box?" -> ""
        "What was the puppy playing with?" -> ""
        "How are the puppy and the boy the same?" -> ""
        "In the movie, who is missing a leg? The boy, the puppy, both the boy and the puppy,
        or no one?" ->  "Both the boy and the puppy."
        "How do you think the puppy was feeling? (after watching first clip)" -> "Happy."
        "How do you think the boy was feeling? (after watching first clip)" -> "Happy."
        "And how did you feel while you were watching that part? (after watching first clip)"
        -> "Good."
        "How do you think the puppy was feeling? (after watching second clip)" -> ""
        "How do you think the boy was feeling? (after watching second clip)" -> ""
        "And how did you feel while you were watching that part? (after watching second clip)"
        -> ""
        "How do you think the puppy was feeling? (after watching third clip)" -> ""
        "How do you think the boy was feeling? (after watching third clip)" -> ""
        "And how did you feel while you were watching that part? (after watching third clip)"
        -> ""
        "How do you think the puppy was feeling? (after watching fourth clip)" -> "Happy."
        "How do you think the boy was feeling? (after watching fourth clip)" -> "Happy."
        "And how did you feel while you were watching that part? (after watching fourth clip)"
        -> "Happy."
        "Rating" -> "0.1"

Example 3:

    (example 3 transcript):
        Interviewer: Have you seen that last movie before, the puppy one?
        Subject: I don't think I have.
        Interviewer: Can you try and tell me what happens in the movie?
        Interviewer: The full beginning, middle, and end.
        Subject: Um, the beginning, the beginning of the movie is, um, so it's the middle,
                 mom gives him the puppy, but he doesn't like it on first because it has three
                 legs, and then towards the end,
        Subject: The last time I played with them.
        Subject: I don't know about it.
        Subject: All right, we both have the same disability.
        Subject: Okay.
        Subject: Didn't go home.
        Subject: How deep?
        Interviewer: What was the boy doing before he got the box?
        Interviewer: What was the puppy playing with?
        Interviewer: How are the puppy and the boy the same?
        Interviewer: In the movie, who's missing a leg?
        Interviewer: The boy, the puppy, both the boy and the puppy or no one?
```

```
        Interviewer: Okay, now we're going to watch a short clip and then we'll talk about it.
        Interviewer: How's the puppy feeling here?
        Interviewer: How's the boy feeling?
        Interviewer: And how do you feel when you watch his part?
        Interviewer: How's the puppy feeling here?
        Interviewer: How's the boy feeling?
        Interviewer: And how do you feel when you watch this part?
        Interviewer: How's the puppy feeling here?
        Interviewer: How's the boy feeling?
        Interviewer: How do you feel when you watch his fart?
        Interviewer: How's the puppy feeling here?
        Interviewer: How's the boy feeling?
        Interviewer: And how do you feeling off that part?
        Subject: Good.
        Interviewer: Okay, we're all done.

    (example 3 output):
        "I hope you enjoyed the last movie. Have you seen it before?" -> "I don't think I have."
        "Can you tell me what happened in the movie? Try to tell the whole story. Remember
        that stories have a beginning, things that happen, and an ending." -> "Um, the
        beginning, the beginning of the movie is, um, so it's the middle, mom gives him the
        puppy, but he doesn't like it on first because it has three legs, and then towards
        the end, The last time I played with them. I don't know about it. All right, we both
        have the same disability. Okay. Didn't go home. How deep?"
        "Do you remember anything else from the story?" -> ""
        "What are some of the things you liked about the movie?" -> ""
        "What are some of the things you didn't like about the movie?" -> ""
        "Who gave the boy a box" -> ""
        "What was in the box?" -> ""
        "What was the boy doing before he got the box?" -> ""
        "What was the puppy playing with?" -> ""
        "How are the puppy and the boy the same?" -> ""
        "In the movie, who is missing a leg? The boy, the puppy, both the boy and the puppy,
        or no one?" ->  ""
        "How do you think the puppy was feeling? (after watching first clip)" -> ""
        "How do you think the boy was feeling? (after watching first clip)" -> ""
        "And how did you feel while you were watching that part? (after watching first clip)"
        -> ""
        "How do you think the puppy was feeling? (after watching second clip)" -> ""
        "How do you think the boy was feeling? (after watching second clip)" -> ""
        "And how did you feel while you were watching that part? (after watching second clip)"
        -> ""
        "How do you think the puppy was feeling? (after watching third clip)" -> ""
        "How do you think the boy was feeling? (after watching third clip)" -> ""
        "And how did you feel while you were watching that part? (after watching third clip)"
        -> ""
        "How do you think the puppy was feeling? (after watching fourth clip)" -> ""
        "How do you think the boy was feeling? (after watching fourth clip)" -> ""
        "And how did you feel while you were watching that part? (after watching fourth clip)"
        -> "Good."
        "Rating" -> "0"

Example 4:

    (example 4 transcript):
        Interviewer: I hope you enjoyed the last movie.
        Interviewer: Have you seen it before?
        Subject: Yeah.
```

```
Subject: The puppy cartoon?
Subject: Yeah.
Interviewer: Can you tell me what happened in the movie and try to tell the whole story?
Interviewer: Remember that stories have a beginning, things that happen, and an ending.
Subject: So first there was a boy that was playing a game.
Subject: So when his mother came in the door was a box.
Subject: She put it down and then a puppy was in the box.
Subject: But the boy didn't want to play.
Subject: Instead he wanted to play on his TV.
Subject: So first he just kicked the puppy, so then he just came back and he did it again.
Subject: So then he just went crazy and then got a ball.
Subject: And then he wanted to play, but then the boy just kicked the ball in the box
         and the puppy ran with it.
Subject: And then
Subject: he dropped the ball back.
Subject: And then the boy stopped when he was playing, and then he went to lose the puppy.
Interviewer: Do you remember anything else from the story?
Subject: That's all I can remember.
Interviewer: What are some of the things you liked about the movie?
Subject: When you like
Subject: And then all of the puppies ran with it and then he got trapped in the box.
Subject: When he kicked the puppy.
Subject: His mom.
Subject: A puppy.
Subject: Playing on his TV.
Subject: A ball.
Interviewer: How are the puppy and the boy the same?
Subject: They're both playing with them.
Interviewer: In the movie, who is missing a leg?
Interviewer: The boy, the puppy, both the boy and the puppy, or no one?
Subject: The boy and the puppy.
Interviewer: Let's watch a short clip from the movie, and then we'll talk about it.
Interviewer: Give me one second, bud.
Subject: Can we do an interview on it?
Interviewer: This is the interview.
Interviewer: Okay, here we go.
Interviewer: You can sit back.
Interviewer: Can you sit in your chair?
Interviewer: How was the puppy feeling?
Subject: Happy.
Interviewer: How was the boy feeling?
Subject: Happy.
Interviewer: How did you feel when you watched that part?
Subject: Happy.
Subject: Ugh.
clip: You've just got to be kidding me.
Interviewer: How was the puppy feeling?
Subject: He was feeling bad.
Interviewer: How was the boy feeling?
Subject: He was feeling, um, angry.
Interviewer: How did you feel when you watched this part?
Subject: Bad.
Subject: Sad.
Subject: Angry.
Subject: Boys.
Subject: Something.
Subject: Get the boy and the puppy.
Subject: I miss the boy.
```

```
        Interviewer: How's the puppy feeling?
        Subject: Very happy.
        Interviewer: How's the boy feeling?
        Subject: Super happy.
        Interviewer: How did you feel while you were watching that part?
        Subject: Happy.
        Interviewer: That's all, thank you.

    (example 4 output):
        "I hope you enjoyed the last movie. Have you seen it before?" -> "Yeah, The puppy
        cartoon? Yeah."
        "Can you tell me what happened in the movie? Try to tell the whole story. Remember
        that stories have a beginning, things that happen, and an ending." -> "So first there
        was a boy that was playing a game. So when his mother came in the door was a box.
        She put it down and then a puppy was in the box. But the boy didn't want to play.
        Instead he wanted to play on his TV. So first he just kicked the puppy, so then he
        just came back and he did it again. So then he just went crazy and then got a ball.
        And then he wanted to play, but then the boy just kicked the ball in the box and the
        puppy ran with it. And then he dropped the ball back. And then the boy stopped when
        he was playing, and then he went to lose the puppy."
        "Do you remember anything else from the story?" -> "That's all I can remember."
        "What are some of the things you liked about the movie?" -> "When you like. And then
        all of the puppies ran with it and then he got trapped in the box."
        "What are some of the things you didn't like about the movie?" -> "When he kicked
        the puppy."
        "Who gave the boy a box" -> "His mom"
        "What was in the box?" -> "A puppy"
        "What was the boy doing before he got the box?" -> ""Playing on his TV."
        "What was the puppy playing with?" -> "A ball."
        "How are the puppy and the boy the same?" -> "They're both playing with them."
        "In the movie, who is missing a leg? The boy, the puppy, both the boy and the puppy,
        or no one?" ->  "The boy and the puppy."
        "How do you think the puppy was feeling? (after watching first clip)" -> "Happy."
        "How do you think the boy was feeling? (after watching first clip)" -> "Happy."
        "And how did you feel while you were watching that part? (after watching first clip)"
        -> "Happy. Ugh."
        "How do you think the puppy was feeling? (after watching second clip)" -> "He was
        feeling bad."
        "How do you think the boy was feeling? (after watching second clip)" -> "He was
        feeling, um, angry."
        "And how did you feel while you were watching that part? (after watching second clip)"
        -> "Bad."
        "How do you think the puppy was feeling? (after watching third clip)" -> "Sad."
        "How do you think the boy was feeling? (after watching third clip)" -> "Angry."
        "And how did you feel while you were watching that part? (after watching third clip)"
        -> ""
        "How do you think the puppy was feeling? (after watching fourth clip)" -> "Very happy."
        "How do you think the boy was feeling? (after watching fourth clip)" -> "Super happy."
        "And how did you feel while you were watching that part? (after watching fourth clip)"
        -> "Happy."
        "Rating" -> "0.8"


### This is the transcript you have to process:
<<<transcript>>>
```
